# Nanopore sequencing of native adeno-associated virus (AAV) single-stranded DNA using a transposase-based rapid protocol

**DOI:** 10.1101/2019.12.27.885319

**Authors:** Marco T. Radukic, David Brandt, Markus Haak, Kristian M. Müller, Jörn Kalinowski

## Abstract

Next-generation sequencing of single-stranded DNA (ssDNA) enables transgene characterization of gene-therapy vectors such as adeno-associated virus (AAV), but current library generation uses complicated and potentially biased second-strand synthesis. We report that libraries for nanopore sequencing of ssDNA can be conveniently created without second-strand synthesis using a transposase-based protocol. We show for bacteriophage M13 ssDNA that the MuA transposase has unexpected residual activity on ssDNA, explained in part by transposase action on transient double-stranded hairpins. In case of AAV, library creation is additionally aided by genome hybridization. We demonstrate the power of direct sequencing combined with nanopore long reads by characterizing AAV vector transgenes. Sequencing yielded reads up to full genome length, including GC-rich inverted terminal repeats. Unlike short-read techniques, single reads covered genome-genome and genome-contaminant fusions and other recombination events, while additionally providing information on epigenetic methylation. Single-nucleotide variants across the transgene cassette were revealed and secondary genome packaging signals were readily identified. Moreover, comparison of sequence abundance with qPCR results demonstrated the technique’s future potential for quantification of DNA impurities in AAV vector stocks. The findings promote direct nanopore sequencing as a fast and versatile platform for ssDNA characterization, such as AAV ssDNA in research and clinical settings.

## INTRODUCTION

Next-generation sequencing techniques for single-stranded DNA (ssDNA) gained attention through the rise of the ssDNA adeno-associated virus (AAV) as a gene therapy vector. Recombinant AAV (rAAV) are preferred gene therapy vectors with currently two approved drugs in the United States (Luxturna and Zolgensma). Their clinical success is driven by a low immunogenic profile and extrachromosomal stability of its genomes. AAV vectors are produced in eukaryotic cell culture by plasmid transfection. For production in mammalian HEK-293 cells, the wild-type AAV genome is separated onto two plasmids such that one plasmid carries the AAV genes *rep* and *cap* (pRepCap) and the other carries the gene of interest (GOI) to be packaged into viral capsids (pITR). The GOI is flanked by the AAV inverted terminal repeat (ITR) sequences, which mediate genome replication and packaging. A third plasmid provides adenoviral helper functions (pHelper) (1, 2) and can be combined with pRepCap to one AAV helper plasmid (3). Based on AAV biology, AAV vectors harbour a single-stranded DNA, but this DNA can be designed to be self-complementary (4).

For quality control of AAV vectors in a clinical context, the state of the encapsulated AAV genome must be tightly monitored. Vector genome quality issues arise from falsely packaged contaminating DNA, which was initially identified by Southern hybridization and quantitative PCR methods to be *rep* and *cap* sequences (5) or sequences from the bacterial plasmid backbone (6). These can make up 0.5% to 6.1% of the cargo DNAs of AAV vectors, dependent on the plasmids used for AAV production (7). The same study found that the amount of contaminating DNA is even higher in self-complementary vectors. These contaminants should be avoided, as the transfer of bacterial sequences is linked to inflammatory response and gene silencing (8, 9) and *cap*-positive vectors have been shown to express AAV capsid proteins, potentially leading to an increased immune response to the vector and thereby impeding its efficacy (10).

While DNA probe-based methods enable investigation of known contaminants, they only allow for a partial view of the sample. In search for unbiased methods to assess contaminations, next-generation sequencing (NGS) protocols have been developed. A first approach to AAV single-molecule sequencing relied on the Helicos HeliScope sequencer and identified low levels of contaminating plasmid DNA, but was deemed too expensive for routine quality control (11). An advancement in this field was an Illumina platform-based method for single-stranded AAV vectors (SSV-Seq), which identified—next to the known contaminations—randomly packaged host cell sequences and AAV purification-specific DNA impurities, as well as helper plasmid-derived impurities (12). AAV self-complementary vectors on the other hand are particularly amenable to NGS by the single-molecule real time sequencing (SMRT) approach, which revealed human DNA-vector chimeras, but requires double-stranded substrates (13). Illumina and SMRT, in general, require extensive sample preparation, including second-strand synthesis which can induce sequencing bias and increases hands-on time.

The rapid transposase-based protocol provided by Oxford Nanopore Technologies offers the advantage of amplification-free direct sequencing, thereby simplifying the sample preparation and potentially eliminating additional sources of bias. In regular rapid library generation, double-stranded samples are fragmented by a transposase and adapters are ligated to the sample fragments as part of the transposase reaction. The sample can then be directly used for nanopore sequencing. We report here the application of this convenient protocol for direct ssDNA sequencing of bacteriophage M13 ssDNA (circular) and of AAV ssDNA (linear) with results obtainable within one working day. In addition, we demonstrate possibilities of large-scale virus genome analysis in the context of AAV biology and manufacturing, aided by the long reads of nanopore sequencing technology.

## MATERIAL AND METHODS

### AAV production and ssDNA extraction

AAV vectors were produced by the calcium phosphate triple transfection method in adherent HEK-293 cells grown in TC150 cell culture dishes, DMEM, 10% FCS, 1% penicillin/streptomycin, 37 °C, 5% CO_2_. Cells were co-transfected by plasmids pRepCap (lab reference pZMB0504, a plasmid encoding the replicases of AAV serotype 2 and the capsid of AAV serotype 9) and pHelper (Agilent Technologies, lab reference pZMB0088) which provides AAV adenoviral helper functions, and pITR (lab reference pZMB0347, encoding fluorescence reporter mKate2 under control of a CMV promoter and human growth hormone polyadenylation signal), which provides the GOI to be packaged into viral capsids (Supplementary Figure S12-14), 37,5 µg DNA per dish, molar plasmid ratio 1:1:1. Three days after transfection, cells were harvested by scraping, pelleted (5 min. 3,000 rcf) and lysed by three freeze-thaw cycles (in 2.5 ml buffer per pellet from two dishes, 20mM Tris, 150 mM NaCl, 10 mM MgCl_2,_ pH 7.5, range -80°C to 37°C). The supernatant was processed further (5 min. 3,000 rcf). Free nucleic acids were then digested with 60 units/ml Benzonase Nuclease (Merck) for at least 30 minutes at 37°C. 3-[(3-Cholamidopropyl)dimethylammonio]-1-propanesulfonate hydrate (CHAPS) was added to 0.5% w/v, and incubated 37°C, 30 min. Additionally, AAV from the culture media were precipitated with ammonium sulphate (3.13 g per 10 ml, 30 min. on ice, pelleting 8,300 rcf, 10 min.) and the pellet was resuspended with the cleared lysate. Further contaminants from this solution were precipitated with a second ammonium sulphate precipitation (25% saturation, pelleting 7,700 rcf, 10 min., 4°C). AAV were then pelleted with a third round of ammonium sulphate precipitation of the supernatant from the second round (55% saturation, pelleting 17,000 rcf, 20 min, 4°C). The obtained pellet was dissolved in PBS (137 mM NaCl, 2.6 mM KCl, 10 mM Na_2_HPO_4_, 1,8 mM KH_2_PO_4_, pH 7.4) and the solution was run through a bed of Poros CaptureSelect AAVX (Thermo Scientific) affinity resin at 1 ml/min. The affinity material was washed with PBS containing 0.05% Tween 20 using at least ten times the bed volume. AAV vectors were eluted with 100 mM citric acid at pH 2.0 (adjusted with HCl) and the eluate was immediately brought to a neutral pH with 1M Tris, pH 8.8. AAV vectors were finally rebuffered (100 kDa cut-off Amicon ultrafiltration spin tubes) to PBS containing a total of 180 mM NaCl and 0.005% Pluronic-F68 and stored at -80°C. A typical yield was 10^13^ DNaseI-resistant particles from five TC150 cell culture dishes.

Vector DNA was extracted by capsid disruption and subsequent silica-affinity based DNA purification. For this, the vector stock was first brought to 100 mM guanidine and 50 mM EDTA from a six-fold stock solution (Tris-buffered pH 8.0). Proteinase K (New England Biolabs) was added to a final concentration of 4 units/ml and incubated 37°C, 1 h and 95°C, 20 min. Vector DNA was extracted from this solution by the NucleoSpin Gel Extraction and PCR Cleanup Kit (Macherey Nagel) as per the manufacturer’s instruction.

### Bacteriophage M13 production and ssDNA extraction

M13mp18-phagemid dsDNA and the corresponding ssDNA were purchased from New England Biolabs. M13KO7 helper phage was produced as described in the supplementary methods.

### qPCR analysis

Quantitative PCR analysis was performed on a Roche LightCycler 480 II using the Promega GoTaq qPCR Master Mix. Primers, annealing temperatures and qPCR qualification data are given in the supplementary methods.

### Nanopore sequencing sample preparation

9 µl or up to 400 ng equivalent of the vector DNA and 1 µl fragmentation mix were used for preparation of barcoded libraries using the Oxford Nanopore Rapid Barcoding Kit (RBK-004) and sequenced with R9.4.1 MinION flow cells on the Oxford Nanopore GridION sequencing machine. Non-barcoded libraries were prepared using the RAD-004 kit. For the sequencing of the commercially obtained M13mp18 DNA, 400 ng (according to the manufacturers’ concentration measurements) of each dsDNA and ssDNA were used.

### Data evaluation

Basecalling was carried out using ont-guppy-for-gridion (v3.0.6) with the high accuracy model (dna_r9.4.1_450bps_hac.cfg). Porechop (v0.2.4) was used for adapter-trimming and demultiplexing. Initially reads with read length ≥1,000 bases and average Phred quality score >5 and later read length ≥500 bases and average Phred quality score >10 were kept for downstream analysis. Reads were mapped to the reference sequences using minimap2 (14) (v2.10-r761) with the map-ont preset. Per-base read coverage was calculated using BEDTools genomecov (15) (v2.27.1) separately for both strands. Assignment of reads to the respective subject sequence was done using BLASTn (16) (max_hsps 3). BLASTn results were analyzed using a custom python script, counting high-scoring segment pairs (HSPs) to each subject and in the case of multiple HSPs of a single query making a subject assignment based on the highest bitscore. Detection of CpG methylation was carried out by realignment of the Nanopore raw data against the respective reference sequence using the re-squiggle algorithm of Tombo (v1.5) (17) and subsequent analysis using DeepSignal (18) with standard parameter settings and the supplied CpG model (model.CpG.R9.4_1D.human_hx1.bn17.sn360). Single-nucleotide variants (SNVs) were called using Longshot (v0.3.5) (19), with a strand bias p-value cutoff of 0.01 and a maximum coverage of 500,000.

Determination of transposase insertion sites was carried out using a custom python script. Reads not subjected to adapter trimming with more than 500 nt length were mapped to the reference genome using minimap2 and the map-ont preset, excluding secondary alignments and alignments shorter than 100 nt. For each read, the alignment closest to the read start was selected and the start coordinate on the reference sequence (the end coordinate for mappings against the negative strand) was taken as an estimate for the insertion site. Each estimate was refined by realignment of the read to a set of 31 reference sequences. These references were constructed by taking the transposase adapter sequence (supplementary information) and adding 75 nt of the reference genome sequence with start positions within a 31 nt window around the original transposase insertion site estimate (which gives a total of 31 reference sequences). If the highest scoring realignment was longer than 100 nt and comprised at most 3 gaps and/or insertions in a sequence window of 10 nt around the start of the genomic sequence, it was considered as a transposase insertion site.

Mappings of sequencing reads to reference sequences performed with minimap2 were analyzed and visualized using a custom python script (see availability section). In case of a circular genomic reference, a single sequencing read may produce two non-overlapping alignments against the reference, discontinuous at the end of the linear subject sequence. Two such alignments were joint if the end-spanning distance between them on the subject sequence was less or equal to 100 nt. In these cases, the subject start of the joint alignment becomes greater than its subject end. In case of visualizing reference mappings of reads with trimmed adapter sequences, the lengths of the non-mapping termini were added to the subject start and subject end as if they were matching. The reads were subsequently binned and plotted with regards to their subject start and end positions. For visualization of untrimmed read mappings, the lengths of these overhangs were displayed in individual subplots.

The propensity of nucleotide regions in single stranded genomes to be double-stranded during transposase-based library preparation was estimated by calculating ss-counts, where a ss-count is the number of times a base is single stranded in a group of predicted foldings. Calculations were based on 100 predicted ssDNA folding structures using Mfold (v3.6) (20) with parameter settings “W=10”, “T=25” and “LC=circular” in case of circular genomes.

## RESULTS

The initial intention of this study was to find a general and fast protocol for AAV ssDNA genome sequencing for quality control of virus-vector batches. We chose nanopore sequencing and reasoned that a convenient way of library creation would be a transposase-based protocol, in which a transposase randomly cleaves the DNA and ligates the fragments to sequencing adapters. If desired, DNA barcodes for sample assignment in multiplexed sequencing could be added. Tagmentation with Tn*5* transposase has been used for Illumina dye sequencing of randomly primed AAV ssDNA before (21). However, the previous approach relied on a multi-step sample preparation to obtain a double-stranded tagmentation substrate. Direct adapter ligation is therefore desirable. Transposases used for rapid library creation in next-generation sequencing are, to the best of our knowledge, not known to use single-stranded DNA (ssDNA) as transposition substrate. We considered methods to obtain double-stranded substrates such as priming the genome at the inverted terminal repeats and subsequently generating the complementary strand with a polymerase. On the other hand, we assume that the AAV inverted terminal repeat (ITR) sequences located at both AAV genome termini are probably already present as dsDNA and could suffice for transposase fragmentation and adapter ligation. Furthermore, AAVs package either DNA strands of their genome with equal probability, with the minor exception of some ITR-modified variants that package only a single-polarity genome (22). DNA extracted from AAV vector stocks might therefore already be in a partly double-stranded state, which should enable direct library creation without prior second-strand synthesis. We followed the two routes of either ITR priming and second-strand synthesis or direct library creation with 10^11^ vector genomes in both cases. Indeed, sequencing reads of comparable quality were obtained from both samples (data not shown). The overall read count in this initial test was low, which we attributed to the low DNA input. Hence, we saw the potential for direct sequencing of AAV ssDNA as a convenient characterization tool. Direct single-molecule sequencing of the ssDNA genome is a preferred method, because hands-on time and thereby additional sources of bias are reduced. We did not observe insertion bias towards the ITRs in this initial experiment and therefore wondered, if ssDNA in general might be a valid substrate for the transposase reaction. At this point of course, we could not rule out strand hybridization as the cause of successful library generation.

### Bacteriophage M13 ssDNA is amenable to direct nanopore sequencing

We tested our hypothesis of generalized transposase-based sequencing of ssDNA by sequencing of M13 phage DNA, which is a commonly used ssDNA reference. Unlike AAV, the bacteriophage M13 packages only one circular strand referred to as the (+) strand during propagation. DNA prepared from this phage therefore is uniform and double strand formation is unlikely. We obtained commercial M13mp18 ssDNA and corresponding dsDNA phagemids. The direct preparation of the transposase-based library from M13 ssDNA and nanopore sequencing was then carried out as before without prior second-strand synthesis. Conforming with our hypothesis, the M13 ssDNA sample was readily sequenced. Sequencing yielded 5,841 reads with an N50 of 6,887 bp for the single-stranded M13 DNA. 5,704 of the total 5,841 reads mapped to the reference sequence of M13mp18. Thereof, 5,591 reads mapped to the (+) strand and 113 reads mapped to the (-) strand, which corresponds to a ssDNA purity of 98%. Furthermore, 3,165 (+) reads and 42 (-) reads passed the filtering criteria to estimate transposase insertion sites (Figure 1 A).

**Figure 1.**
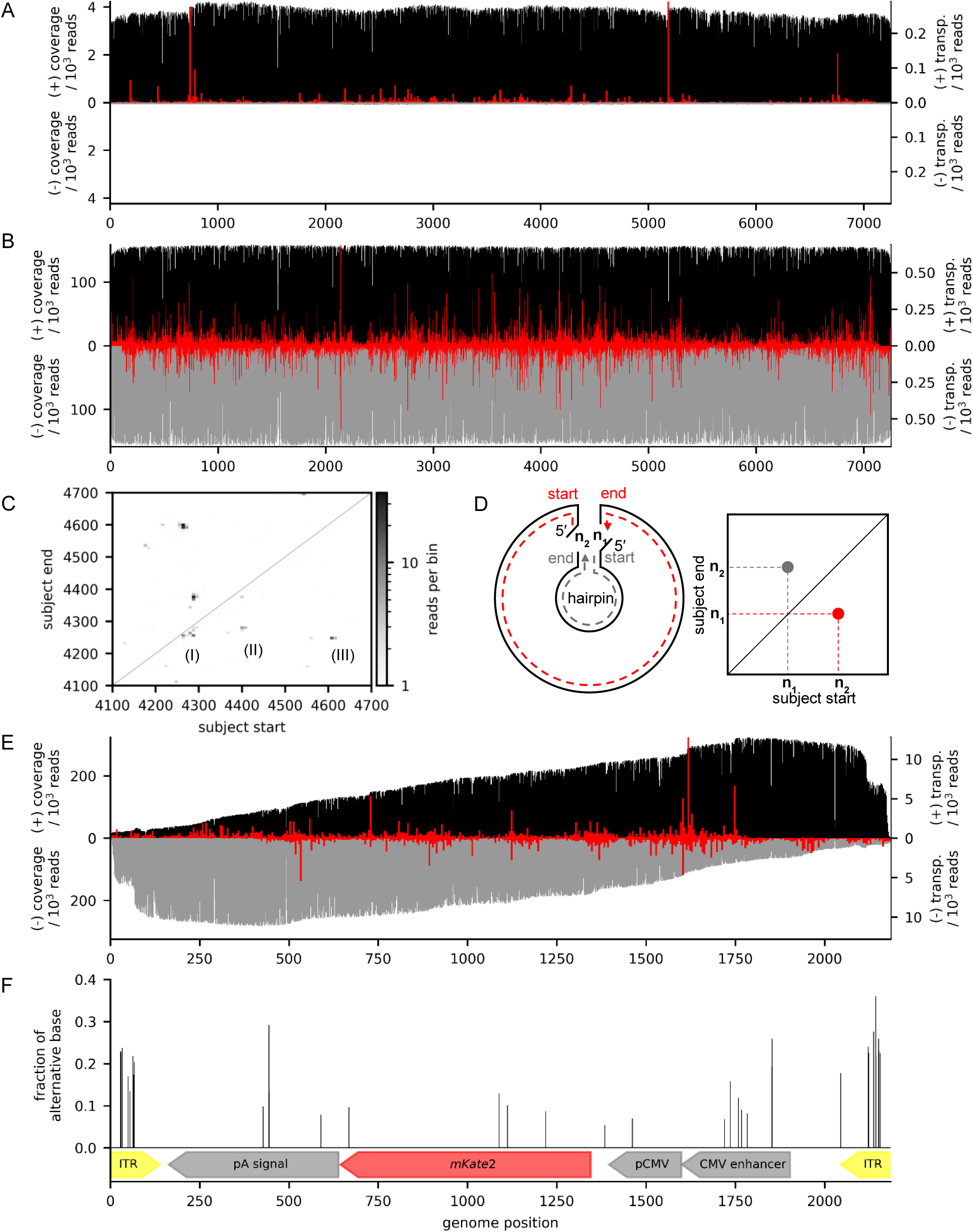
Strand-specific sequence coverage of M13mp18 samples and the rAAV genome, as well as its unveiled single-nucleotide variants and transposase insertion sites. **A**: Sequence coverage for M13mp18 ssDNA (black and grey for (+) and (-) strand) is near-constant, which is expected for a circular molecule and long reads. Overlaid transposition sites (red, 15 nt bin width) reveal that 18% of reads start from one of three prominent starting points. **B**: Coverage for M13mp18 phagemid dsDNA is homogenous throughout the sequence and both strands. Transposase insertion sites (red, 15 nt bin width) had no outstanding hot spots, indicating that the mode of action of the transposase is inherently different between ssDNA and dsDNA. **C**: Plot of binned (bin with 8 nt) subject start positions against binned subject end positions for all mapped adapter trimmed reads of the M13mp18 ssDNA. The non-mapping termini of each read were added to the subject start and end as if they were matching. Two (II and III) of three prominent hot spots (I, II, and III) below the diagonal line have a counterpart above the diagonal line revealing a mirror symmetry. **D**: Illustration on how the symmetry in the subject start-end plot is a result of transposase-induced double strand break within the hairpin, leading to two possible 5′-ends for sequencing adapter ligation (left panel) and subsequently two possible reads, one long read (dashed red line) and one short read comprising the hairpin (dashed grey line). Since the start position of one read should be the end position of the other and *vice versa*, the symmetric pattern emerges from the hairpins in the start-end plot (right panel). **E**: Coverage of the rAAV genome (black and grey for (+) and (-) strand) is constantly increasing towards the 3′-end, as expected for a linear substrate, until it reaches a plateau and suddenly halves within the ITRs. Both strands are covered, as AAV packages both strands during production. Overlaid transposase insertion sites (red, 5 nt bin width) exhibit an even distribution across the genome with few hot spots. The lack of transposase insertion sites towards the ends of the linear sequence is attributable to the applied read length cut-off. Furthermore, a 3 bp discrepancy to the theoretical sequence is uncovered near the 3′ ITR. **F**: Fraction of alternative bases at identified sites of single-nucleotide variants (SNVs).

From the M13mp18 dsDNA sample, 384,091 reads with an N50 of 7,224 bp were generated and in contrast to the ssDNA sample, reads of the phagemid sample mapped to both strands with near-equal distribution. Of 382,079 total mapped reads, 191,446 reads mapped to the (+) strand and 190,633 reads mapped to the (-) strand, respectively. For the estimation of transposase insertion sites, 110,994 (+) reads and 105,783 (-) reads passed the filtering process (Figure 1 B).

We observed reduced sequence quality with the ssDNA samples. Average sequence similarities to the M13mp18 reference sequence were at 93.38 % for M13 dsDNA and 89.88 % for M13 ssDNA on single-read level. Similarities were calculated as weighted average considering varying alignment lengths from over 10,000 BLASTn hits against the reference sequence. It is possible that the helicases, which are located at each pore and are employed for double-strand separation and the reduction of sequencing speed do not function properly with ssDNA substrates.

Regarding reactivity, we found that the ssDNA samples gave a significantly reduced output compared to that of the dsDNA phagemid sample. On first sight, the mapped reads were evenly distributed over the reference sequence for all data sets, however when we plotted the corresponding transposase insertion sites, hot spots were much more apparent for the ssDNA sample (Figure 1 A and B) and 18% of total reads started at these positions. In a first analysis there was no clear correlation of these hot spots to the substrates ss-count in Mfold (20), which would indicate transposase preference towards dsDNA stretches (Supplementary Figure S1).

### The transposase acts on hairpin loops

For a more detailed analysis of transposition insertion sites, we binned subject start positions and end positions for all mapped reads as detailed in the Methods section and plotted subject start bins against subject end bins (Figure 1 C shows an excerpt and Supplementary Figure S2 shows the full plot). Mapped reads of M13 dsDNA showed a nearly even distribution of start and end positions across the plot, indicating that transposase insertion is indeed random on dsDNA. Reads from M13 ssDNA on the other hand accumulated at distinct start-end positions, which fits the transposase insertion site preference seen in Figure 1 A and further shows that, as expected, reads generated from the circular M13 genome end at defined positions upstream of the observed read start. Figure 1 C shows the range 4,100 to 4,700 of the M13 ssDNA reads start and end positions. Interestingly, for many bins below the diagonal line with subject start > subject end there is a corresponding bin mirrored above the diagonal line comprising very short reads with subject start < subject end. This points to the conclusion that transposition on ssDNA, at least to some extent, depends on the formation of hairpin loops. This is because transposase insertion in a hairpin yields one long fragment and one short fragment that contains the hairpin. Both resulting fragments can be sequenced provided that two individual transposase insertion events occur in which sequencing adapters are ligated to either of the two resulting 5′-ends. Since the transposase will always produce a double-strand break within the hairpin loop, the start position of one fragment should always match the end position of the other and vice versa, which results in the mirrored pattern. An illustration of this reasoning is provided in Figure 1 D. Figure 1 C shows three prominent long fragments (I, II, and III) below the diagonal line. Fragments (II) and (III) have a corresponding short fragment in the upper part of the plot, whereas (I) does not have a corresponding fragment. To exemplarily confirm the presence of hairpins in M13 ssDNA, Mfold (20) was used to predict the secondary structure of bases 4,100 to 4,700 (Supplementary Figure S3). For each putative hairpin the results show a predicted intramolecular interaction between two regions that fits the expectation regarding loop size, which can be derived from Figure 1 C. Bins very closely below the diagonal line do not have a corresponding bin above the diagonal line, which means that, either the DNA loop did not yield a long enough fragment to be sequenced or the transposase also truly acts on single-stranded DNA. We repeated the experiment with M13KO7 helper phage propagated in our lab and obtained a similar result (Supplementary Figure S4). The patent literature suggests that the transposase in the Oxford Nanopore protocol is the MuA transposase (23). The mechanism of this transposase is complex, and ssDNA has been shown to be a cleaved substrate to generate what was termed “Mu-ends” (24), but not a target for a transposition event. Although our findings point to the transposase acting on hairpin loops resulting from self-annealing on short stretches of a few bases, it is unclear, whether there is additional activity on true ssDNA. For AAV vector quality control by direct library generation and nanopore sequencing and direct sequencing of ssDNA in general, these results mean that the presence of both strands in the sample is not a necessity and that also true ssDNA is accessible by this method.

### Nanopore ssDNA sequencing allows for direct, amplification-free sequencing of AAV vectors

As we had observed a relatively low AAV read count in our initial test, we next improved ssDNA extraction from AAV to gain more reads. In the end, we settled with an AAV purification protocol based on Benzonase nuclease digest of the producer cell lysate, ammonium sulphate precipitation and subsequent Poros Capture Select AAVX affinity chromatography. Inactivation of residual Benzonase and capsid disruption for ssDNA release was then performed with 50 mM EDTA, 100 mM guanidine and proteinase K at slightly basic pH. Afterwards, ssDNA purification from this solution was achieved by silica-adsorption chromatography with a commercial kit and the eluate was used for the transposase reaction. Using this protocol, we performed two independent sample preparations with a time delay in between of three months starting from individual cryo-cultures of producer cells, with five TC150 cell culture dishes each. These samples will be referred to hereafter as sample 1 and sample 2 with their sequencing runs being run 1 and run 2. We obtained about 10^13^ DNase I-resistant particles after affinity chromatography from both cultures. From these two AAV preparations we were able to obtain 50 µl DNA solutions with optical densities of OD_260, 10mm_ = 0.40 and OD_260, 10mm_ = 0.85, corresponding to an equivalent total of about 1.0 µg and 2.1 µg dsDNA. Since DNA in these samples might be partially single-stranded and double-stranded, and since these forms have different absorption coefficients at 260 nm, we prefer to use volumes and optical densities for indications of quantities. Agarose gel electrophoresis of these two samples, directly after extraction and after sample freeze-thaw, showed distinct bands attributable to the rAAV genome in single-stranded, hybridized and aggregated states (Supplementary Figure S5).

We used 9 µl of each of the two AAV samples for the transposase reaction and sequenced sample 1 on a pre-used flow cell with sample assignment by barcodes (run 1). A fresh flow cell was used for sample 2 without barcoding (run 2). Again, as expected, both samples were readily sequenced, but gave vastly different read counts that passed our initial length quality threshold of ≥1,000 bases read length (22,174 reads for run 1 versus 291,036 reads for run 2). We performed a first mapping analysis of these reads and found that the vast majority of these raw reads mapped to the reference genome (Table 1 A). Coverage steadily increased until it reached a stable plateau at about half the genome length. At the ITRs however, a sudden decrease in coverage was observed (Figure 1 E displays run 2). However, we found that many mappings that end in the ITRs are stemming from reads that have an unmapped 3′ extension of lower-quality bases, which can be attributed to the remaining ITR (Supplementary Figure S2). Nonetheless, ITR coverage is still 222,712-fold in run 2 and ITR sequences are thus accessible by nanopore sequencing despite their known tendency to form secondary structures. In the transposase insertion site analysis of reads longer than 500 nt of AAV sample 2, 220,075 (+) and 189,481 (-) reads passed filtering. Strand-specific hot spots were again apparent, although overall, most reads started throughout the genome (Figure 1 E). These hot spots again did not correlate with the DNA fold and correlations to the GC content were minimal (Supplementary Figure S6). Interestingly, the read start pattern seems to be a combination of the patterns observed for M13 ssDNA and dsDNA.

**Table 1.**
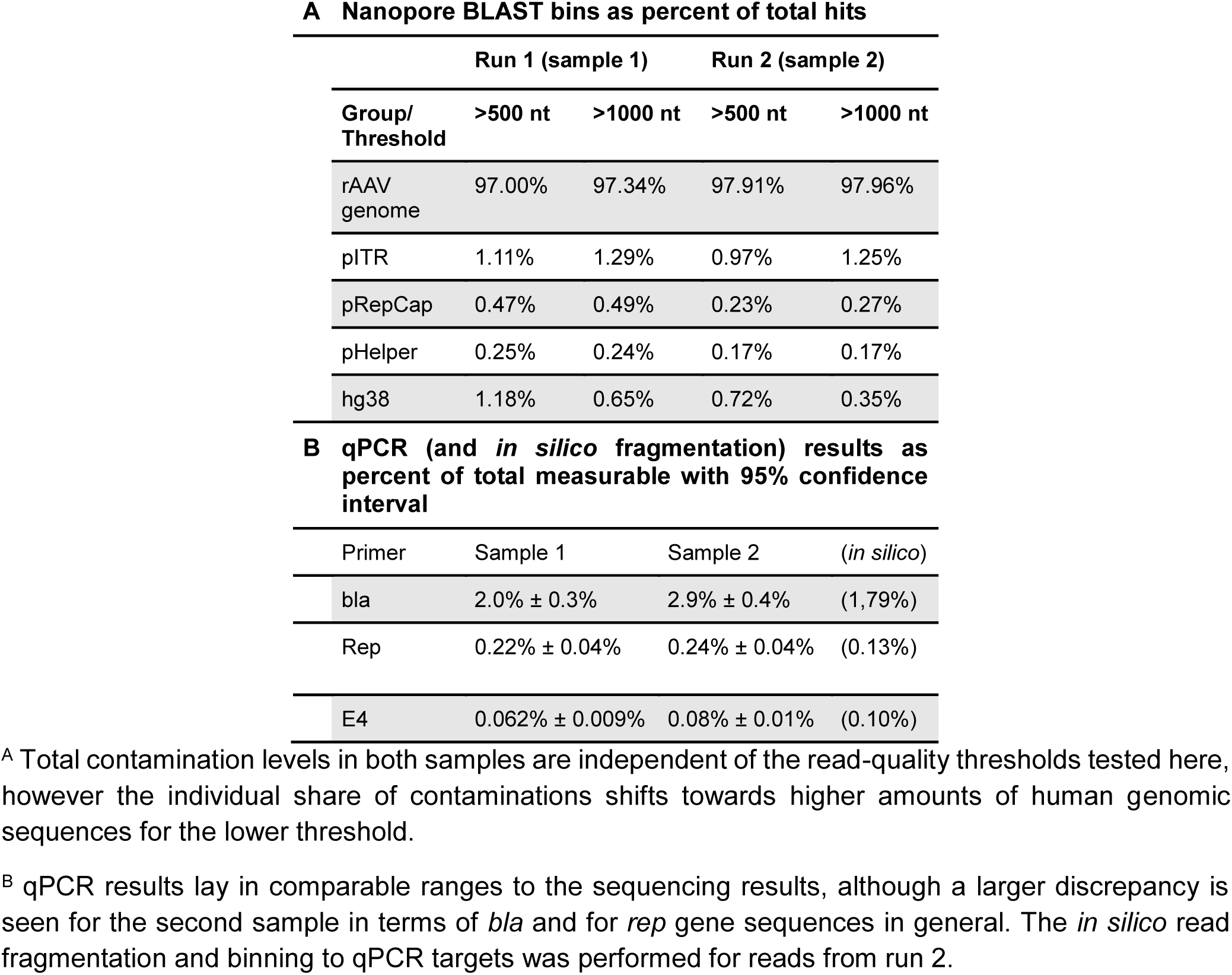
BLASTn read assignments and qPCR results for two independently produced and sequenced rAAV samples (sample 1 and 2).

### Direct nanopore AAV ssDNA sequencing reveals single-base heterogeneity and methylation status

Comparison of the assembled genome to the reference sequence revealed single-nucleotide variants, as seen before for rAAV (12). The high coverage enabled these conclusions despite the relatively low base accuracy of about 93%, which is an intrinsic property of the current nanopore sequencing technology. SNVs were located within ITRs in the short hairpins (B-C internal palindromes according to common nomenclature) and were transversions as well as transitions with an individual abundance of about 20%. We were able to link ITR SNVs to the two possible ITR configurations in FLIP and FLOP orientation, so that the found ITR SNVs are in the end expected to arise. On the other hand, prominent SNVs across the transgene cassette were mostly transitions with a hot spot located in the polyadenylation signal and throughout the CMV promoter with an abundance up to 30% (Figure 1 F). Only two and zero variants could be called for ssDNA M13 and dsDNA M13 samples, respectively. Raw reads were also analyzed for methylated CpG sites separately for both strands using a custom software workflow with Tombo and DeepSignal (18). The studied rAAV genome as encoded on the pITR plasmid contains 129 CG dinucleotides, 123 of which have been mapped in run 2. When we compared reads from run 2 to reads of *in vitro* amplified rAAV genomes, no substantial methylation was identified. However, these results are based on the current algorithms used by the applied software and need to be verified by additional experiments.

### Qualitative statements on the abundancy of impurities are possible

Our nucleotide reference database so far contained only the rAAV genome. We next extended this database to include the human genome build hg38 (GCF_000001405.39) as well as the utilized producer plasmids. Now, of all reads that passed the length quality threshold of ≥1,000 bases, 96.73% (run 1) and 99.92% (run 2) gave a BLASTn-hit with our database. Of the reads not assigned to our database, 17% (101 reads, run 1) and 13% (33 reads, run 2) gave a hit against the National Center for Biotechnology Information (NCBI) Nucleotide database. We performed further analysis only with reads assigned to our database. The assignments calculated as percent-contaminants are summarized in Table 1 A. Of the contaminants, pITR sequences showed the highest prevalence, representing 1.29% (run 1) and 1.25% (run 2) of total reads attributable to our database. They were followed by hg38 sequences, representing 0.65% and 0.35% of all reads. The proportion of reads that map to the pRepCap was 0.49% and 0.27% each. 0.24% and 0.17% of attributable reads in both runs were assigned to the adenoviral helper genes. Notably, these results also depended on where the origin of sequence numbering was set on the reference plasmid, which is a limitation of the mapper. We therefore set the numbering origins to 2,000 bp downstream of the f1 ori for all three plasmids.

We performed additional qPCR analysis with primer sets that allow for the amplification of the CMV promoter within the rAAV genome, *rep* gene, *E1* gene of the adenoviral helper plasmid as well as the ampicillin resistance gene *bla*, which is present on all three producer plasmids. Results are given as percentage of the combined absolute copy number of all four measurements (Table 1 B). As expected for the contaminants, the *bla* gene was present with the highest proportion of 2.0% ± 0.3% (95 % confidence interval) in sample 1 and 2.9% ± 0.4% in sample 2. This result can be compared to the nanopore reads that map to one of the three *bla* containing producer plasmids, which had a combined share of 2.0% (run 1) and 1.7% (run 2) of all referenced reads. The qPCR result for *bla* thereby matches very well for run 1 but is slightly higher than expected from the nanopore analysis for run 2. On the other hand, the proportion of *rep* genes found by qPCR and sequencing matched well for run 2 but were off by a factor of two for run 1. Furthermore, adenoviral helper genes were underrepresented by qPCR compared to the sequencing results in both samples.

We next wondered, whether certain contaminations are present in the capsid as small fragments below 1,000 nt and if our initial length quality threshold of ≥1,000 nt would cause the deviations between qPCR and nanopore sequencing. We therefore re-analyzed our datasets and included all reads above 500 nt and by doing this, the accepted read count for run 2 increased from 291,036 reads to 647,810 reads. The results of this analysis showed that, while the overall share of contaminants within the sample stays roughly the same regardless of the thresholds tested, the share of individual contaminants shifts. We found that the proportion of plasmid-derived contaminants remained mostly constant for both analyses, whereas the proportion of human genomic contaminants doubles for the lower threshold.

Since the BLASTn binning could be distorted compared to qPCR analysis, because long reads might span fusions that contain two or more qPCR target sites, we performed *in silico* read fragmentation to better compare nanopore sequencing and qPCR data. We subsequently split all reads from run 2 (> 500 nt) into fragments of 500 bp and removed resulting fragments below 300 bp. Subjecting a subset of 100k read fragments to the same BLASTn analysis as conducted above gives slightly different results, as the fraction of reads attributed to the rAAV genome drops to 96.42 %. Accordingly, pITR and hg38 contaminants rise to 1.35 % and 1.55 %, respectively. pHelper and pRepCap contaminants account for 0.47 % and 0.21 %, respectively. This leads to a total of 2.03 % of reads attributed to one of the bla containing producer plasmids in sample 2 after *in silico* read fragmentation. Another possibility is to bin the fragmented reads to a reference library that only contains the qPCR target sequences for direct comparison (CMV promoter, *rep, bla*, E4 gene). Here, the share of E4 sequences is comparable between the methods, while *bla* and *rep* sequences are slightly underrepresented (Table 1 B). Clearly, at this point, a more descriptive data evaluation tool is needed to find the source of this disparity.

### Direct sequencing reveals the molecular state of the genome and its contaminants

As we use a direct sequencing approach, each read represents a single 3′-end ssDNA fragment of a natively packaged nucleic acid, presuming that it was fragmented only once by a transposase. This makes the fragments’ GC content a calculatable (from the known sequence) as well as measurable (from sequencing) quantity for a given fragment length, at least for the recombinant AAV genome. Conclusions on the molecular state of the genome and its contaminants can then be drawn from a %GC *versus* read length plot showing reads selected based on the BLAST assignment, as shown in Figure 2. In these plots, when assuming no sequence preference of the transposase, reads of originally circular molecules ideally appear as single points, as these have a constant %GC content and the same read length, independent of the cut site (for single cut genomes). Accordingly, reads of linear fragments will produce a vertical line (|), if the GC content is constant along the DNA and a slanting line (/ or \), if the GC content increases towards one end. The latter fragment will result in an upper-case lambda (Λ) structure when both strands are present, because both directions are sequenced. In such a plot and according to expectations, the M13 reads group around the theoretical GC content and length with conical tailing towards shorter reads (Supplementary Figure S7). We suspect the latter to arise from double transposase fragmentations and premature sequencing breakoffs. In the AAV sequencing runs on the other hand, reads that gave a BLAST hit with the rAAV genome showed a more complex mirrored lower-case lambda-like pattern. The pattern becomes easier to spot in the large data volume when reads are displayed in a heatmap, as shown in Figure 2 A for run 2 (refer to Supplementary Figure S8 for run 1). The read-length histogram underlaid in Figure 2 B for the same data set further shows that most read lengths are at or below the theoretical genome length, which is 2.2 kb. The shape of the data distribution in the plot is a function of the fragment’s nucleic acid composition. We therefore simulated the transposase reaction for the rAAV genome and found that in the plot the measured data are shifted slightly towards lower GC content compared to that of the simulation (Figure 2 A, green line and supplementary information for the simulation script). Single-read analysis showed that the GC-rich 3′ ITR is often sequenced incompletely, which explains the shift.

**Figure 2.**
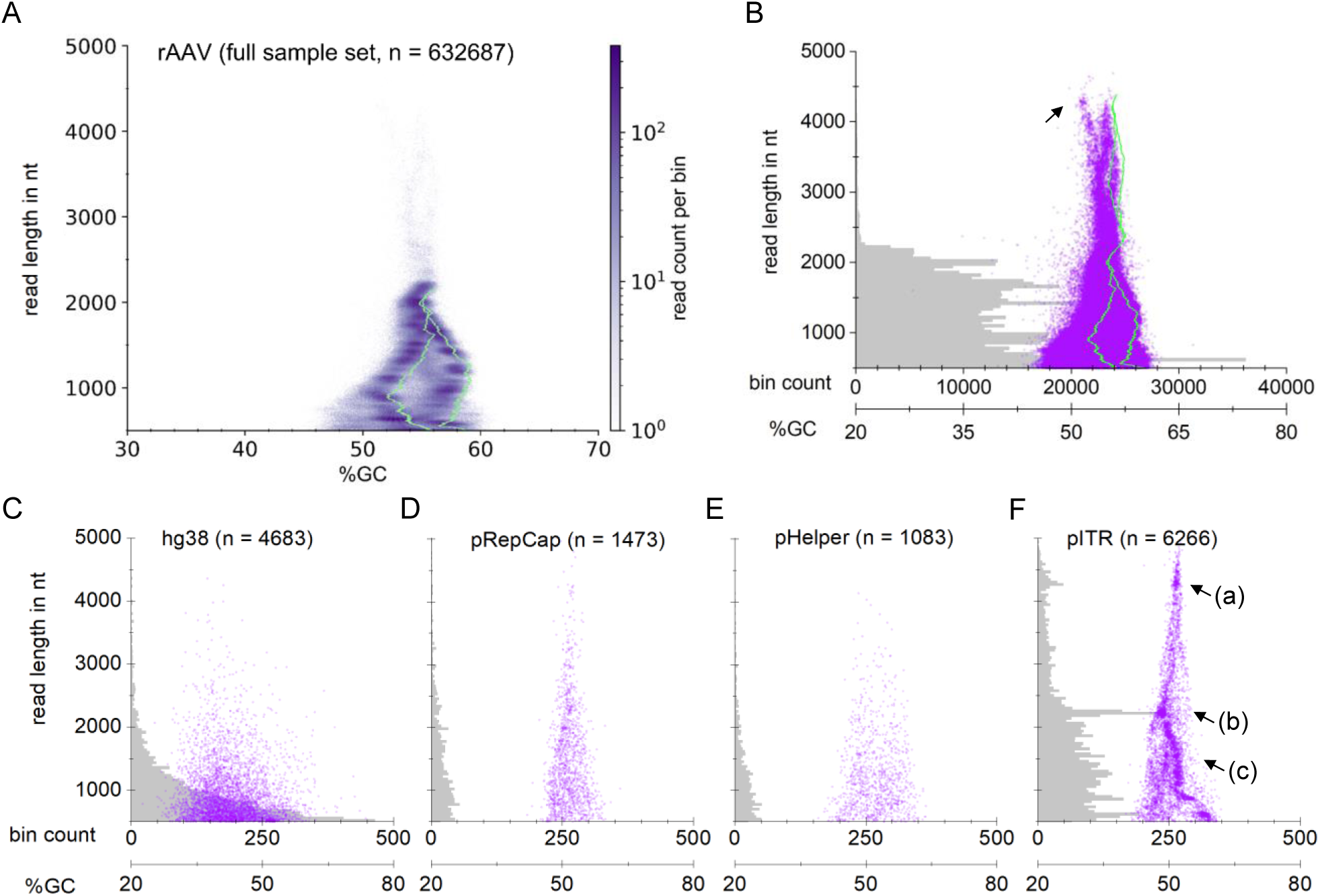
Molecular state of the recombinant AAV genome and its nucleic acid contaminants in %GC vs. read length plots. Grouping by BLAST assignments for run 2. **A**: A heatmap of reads in the rAAV bin (2D histogram with hexagonal bins and logarithmic scale) reveals a distinct underlying structure. The structure is a function of the genome’s nucleic acid sequence and can be predicted by simulating one transposase fragmentation reaction per genome (green line). The data set is shifted towards lower %GC compared to the simulation, because many reads miss part of the 3′-ITR sequence. **B-F**: Each translucent purple dot represents one individual read. A histogram of read length distribution is underlaid in grey (histogram bin size of 40 nt). **B**: Display of the full sample set in the rAAV genome bin. The histogram illustrates that most reads are genome monomers. The dot plot reveals that larger forms of the genome are packaged in the capsid as well. The simulation (green line) and single-read investigation unveil these as covalent genome head-to-tail fusions. The black arrow indicates genome-backbone fusions (see text). **C**: Reads in the hg38 bin show no pattern in their %GC content and an exponentially decaying size distribution, indicating packaging of fully random fragments in favour of shorter ones. **D, E**: Reads in the pRepCap and pHelper bin show no distinct structure, however the prevalence of longer fragments as opposed to the size distribution of hg38 reads hints on a different mechanism of packaging. **F**: pITR binned reads appear with all possible lengths within the AAV packaging limit and show distinct structures, indicating non-random packaging. The arrow indicates a hot spot of reads starting in the bacterial *ori*, spanning the transgene cassette and terminating in an ITR. The arrow (b) indicates a hot spot of reads having the same read starts as the fragments of (a) but terminating at the adjacent ITR. The arrow (c) indicates fragments of the reads of (b), all terminating in the adjacent ITR.

Hot spots of transposition sites are also apparent in this plot in accordance with Figure 1 E. Notably, a larger proportion of AAV-assigned reads are shorter than 1,000 nt, which hints on double fragmentations. Plotting all 632,687 mapped reads directly reveals additional reads of a distinct distribution, which are longer than the theoretical rAAV genome (Figure 2 B), although of all reads in the rAAV genome bin, only 0.9% are longer than 2,300 nt. We reasoned that these could be genome head-to-tail (or tail-to-head) fusions, where a head-to-tail fusion is a concatemer of two genomes fused over a junction ITR so that the terminator of one genome is adjacent to the promoter of the next genome. Indeed, a simulation of such a fusion (green line, Figure 2 B) matches well with the oversized reads, except for a protruding point cloud at about 4 kb read length (Figure 2 B black arrow), which are genome-backbone fusions as determined by later single-read investigation. Similar plots for the other BLAST bins reveal differing molecular states of the individual contaminants. Reads assigned to hg38 appear to be of completely random human origin, with an exponentially decaying size distribution (Figure 2 C). Reads assigned to pRepCap at first sight also appear to be randomly fragmented and packaged as indicated by the overall triangle shape of the point cloud (Figure 2 D). However, the underlying size distribution does not show an exponential decay of read abundancy with read length and this could hint on a different mechanism of packaging compared to packaging of human genomic sequences. The same can be observed for pHelper-derived sequences (Figure 2 E). In contrast, the distribution of reads assigned to the pITR backbone appears to follow clear rules (Figure 2 F) and three features of this bin come immediately to attention as labeled in Figure 2 F: (a) An accumulation of reads is located at around 4.25 kb read length, another accumulation at around 2.25 kb read length, as also evident from the underlying histogram plot and (c) a slanting cloud of data points spanning from the shorter length accumulation all the way to the lower limit of read length. Accumulations in this plot can indicate transposase insertion bias on linear DNA resulting in fragments of similar length, although the prevalence of this 2.2 kb point cloud is more prominent than other hot spots for the rAAV genome. Another explanation might be a circular fragment. Conveniently, individual read subsets can also be investigated on single-read level as further elaborated below.

### Single-read investigation confirms the global analysis and highlights the system’s packaging flexibility and susceptibility for recombination

We took the opportunity long-read sequencing data hold and investigated reads of interesting appearance individually by annotating producer plasmid features (PlasMapper tool of Geneious R9 software) on each sequencing read. We started analysis with a subset of 200 randomly chosen reads from the rAAV bin of run 2 and of these, 198 reads (99%) were full-size transgenes or fragments of it, present in both polarities as exemplarily depicted in Figure 3 A. Of these, one (0.5%) was a head-to-tail fusion as depicted in Figure 3 B. Two reads (1%) were transgene-backbone fusions with one junction ITR (Figure 3 C). Furthermore, we were able to annotate 3′-terminal ITR sequences (similarity threshold: 65%) for 149 reads of the subset (74.5%). This is especially interesting, because AAV packages its genome 3′ to 5′, whereas the sequencing direction is 5′ to 3′. This means that we can directly sequence the packaging signal and it also provides a measure on double fragmentations. We repeated the analysis for all 647,810 reads longer than 500 nt using BLAST (E-Value-cutoff of 1e-5) and came to a very similar result of 493,916 reads (76%) with 3′-terminal ITR sequences. We next expanded the subset of the rAAV bin to 10,000 randomly selected reads, and of these, 98 reads (1%) were longer than 2,500 nt. 81 reads (83%) of these overlong genomes terminated in an ITR sequence. 75 reads (76.5%) were rAAV genome-genome fusions. The remaining reads were genome-backbone fusions with a 3′ transgene (except for one read) and these findings were in good agreement with our %GC *vs*. read length plots and simulations while also providing indications on the abundances of fusions. We can extrapolate from these numbers the total content of genome monomers in our sample to be 96.95% (abundancy of monomers multiplied with rAAV bin size), although notably, this extrapolation is only valid under the assumption of quantitative sample preparation, no transposase insertion bias and equally likely packaging of both polarity genomes.

**Figure 3.**
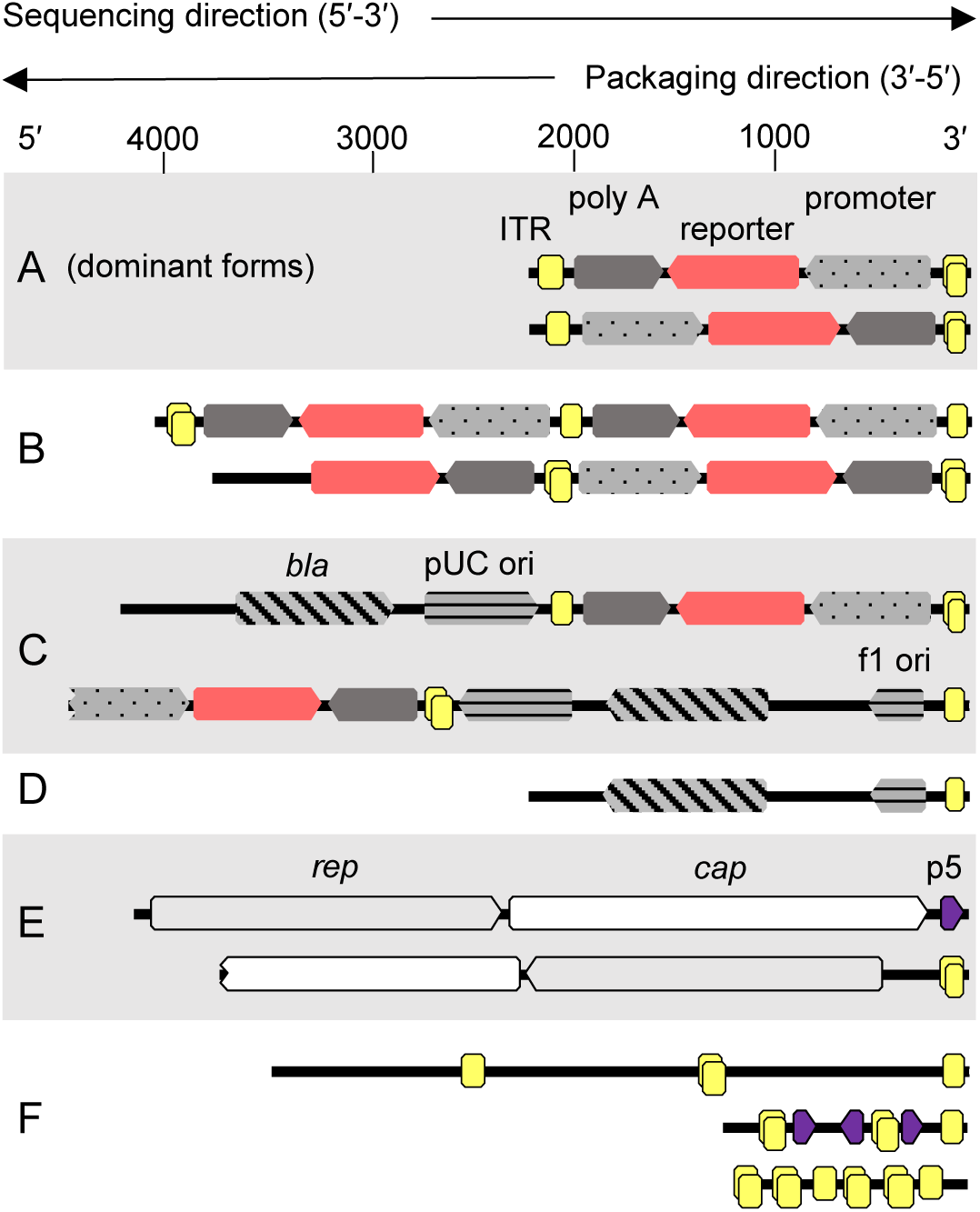
Read examples from sequencing run 2 with features annotated with Geneious (similarity threshold: 50% for ITRs and 65% for other features to account for features that are only partially present). Owing to their self-complementary sequence, ITRs may be double-annotated (two overlaid yellow boxes). See text for qualitative statements on sequence abundancies **A:** Transgene of both polarities with ITRs (boxed yellow), CMV promoter (dotted grey), and terminator (dark grey). **B:** Transgene head-to-tail or tail-to-head fusions of both polarities with a single junction ITR. **C:** Transgene-backbone fusions with bacterial origins of replication (horizontally striped grey) and *bla* gene (diagonally striped grey). **D:** Predominant 2.2 kb backbone sequence found in the pITR bin. **E:** *rep* (boxed light grey) and *cap* (boxed white) sequences with 3′ p5 promoter (boxed dark purple) or ITR sequences. **F:** ITR-ITR or ITR-p5 fusions found in sequences rejected during binning. The CMV promoter can be annotated between the ITRs of the upper read with 35% similarity.

Next, we were interested in the junction ITRs between two fused genomes and genome-backbone fusions, to eventually find hints on their origins. We previously described that in our pITR plasmid, just as in many other available ITR plasmids, the ITR adjacent to the transgene’s polyadenylation signal harbours an 11 bp deletion within the B-C hairpins (25). In addition, the ITR complementary A-region is shorter in both ITRs compared to the ITR reference sequence. Mapping of genome-backbone fusions (n = 12) to our plasmid reference showed identity (including the deletion) to the plasmid reference within the limitations of current nanopore sequencing technology (Supplementary Figure S9). Junction ITRs between two fused genomes on the other hand were much more diverse. We constructed a reference sequence with two genomes fused by a complete ITR including two D-sequences, either in FLIP or in FLOP orientation, and mapped fusions to this reference with respect to their orientation. It became immediately apparent that, independent of genome orientation, the mapping quality dropped downstream of the B-C hairpins, which either hints on difficulties in sequencing, error prone recombination at this position or both (Supplementary Figure S10). Interestingly, of all manually investigated junctions (n = 41), only two (5%) clearly harboured the 11 bp deletion, although the evaluation on single-read level was complicated by the overall elevated number of mismatches in this region compared to our reference. The 3′-terminal ITRs of the same reads (n = 32) showed a similar deletion only in four cases. Regarding ITR orientation (the ITR internal palindrome B-C can be inverted compared to rest of the ITR due to the genome replication scheme), we found that both FLIP (not inverted) and FLOP (inverted) orientations were present at the junctions and at the 3′ terminal ITRs and that genome orientation was biased towards FLOP orientation when the polyadenylation signal was 5′ of the junction (n = 17 in FLOP and n = 4 in FLIP orientation *versus* n = 6 in FLOP and n = 16 in FLIP when the CMV promoter was 5′ of the junction). Junction ITRs furthermore had two D-sequences.

Further single read investigation of the three regions of interest within the pITR bin (Figure 2 F) showed that reads from the 4.25 kb accumulation (arrow a) were mostly backbone-genome fusions with conserved read starts within the bacterial origin of replication and a full 3′ transgene (n = 13 of an 18 read subset with length threshold 4,200 to 4,300 nt). Reads from the 2.25 kb accumulation (arrow b) were mostly reads starting at the same conserved position within the plasmid’s bacterial origin of replication, but ending in the adjacent 3′ junction ITR between backbone and transgene (example in Figure 3 D; n = 53 of a 75 read subset with length threshold 2,200 to 2,300 nt). Reads from the slanting cloud of data points (arrow c), on the other hand started throughout the bacterial backbone and terminated in an ITR as well. However, these were of the inverse orientation compared to the reads of the described accumulation (n = 62 of a 200 read subset with length threshold 500 to 2,000 nt). To a smaller extent, shorter reads of this orientation also ended in an ITR (n = 37 of the same 200 read subset with length threshold 500 to 2,000 nt).

### The p5 promoter is a secondary packaging signal and is prone to recombination with ITR sequences

Reads from the pRepCap bin often terminated 3′ in a p5 promoter sequence, which lead us to further investigate on the phenomenon (example Figure 3 E top). We searched for 3′-located p5 sequences using BLAST (E-Value-cutoff of 1e-5) and found 527 reads (0.08%), and these had the *cap* coding sequence 5′. Of these, 303 reads had the p5 promoter strictly 3′-terminal and the coverage sharply dropped at position +12 downstream of the promoters TATA-box (coverage at position +11: 71%, +12: 42%, +13: 13%, Supplementary Figure S11). On the other hand, 158 reads had the p5 promoter followed 3′ by a terminal ITR. In 59 of these cases the ITR D-sequence was located directly adjacent to the p5 promoter and in the other cases a maximum of 154 nt lay between p5 and the D-sequence. Long reads in the pRepCap bin also had an ITR without p5 sequences, when the *rep* gene was 3′ (Figure 3 E bottom, n = 3 in a subset of 47 of reads longer than 3 kb). Significant recombination was also evident in the pHelper bin. 41% of reads in the investigated subset of this bin had 3′ ITR sequences (n = 82 in a random 200 read subset), often in combination with CMV promoter sequences (n = 19). ITR sequences were furthermore especially prevalent in reads where binning had failed (560 reads total in run 2). A random subset of 260 reads of these had a mean length of 715 nt and, despite their short length compared to the other bins, 47 reads of the subset (18%) had more than one ITR sequence (read examples shown in Figure 3 F).

## DISCUSSION

Nucleic acid contaminants in AAV vector stocks for gene therapy are gaining attention alongside the increase of therapeutic doses from 10^12^ viral genomes per kg in the first authority approved product Glybera to recently approved 10^14^ viral genomes per kg for Zolgensma, both single-dose systemic applications (26, 27). Potentially, even higher doses in multi-administration therapies, like cancer gene therapy, are conceivable. The United States Food and Drug Administration recommends for a vaccine dose that residual cell-substrate DNA should be ≤10 ng and the median DNA size should be of 200 bp or lower (28). Vector manufacturers take extensive precautionary measures to ensure a homogenous product and preempt tighter AAV-specific regulations. These measures include the use of bacterial backbone-depleted circular supercoiled plasmid-derivatives, which were shown to be effective in significantly reducing false encapsidation of prokaryotic sequences in rAAV (7). Other strategies involve the use of closed-linear derivatives (29) or plasmid insertions of uncritical stuffer DNA beyond the AAVs packaging limit, to avoid packaging of the bacterial backbone and antibiotic-resistance gene (30). Monitoring of contaminants is a routine task in vector manufacturing and new time-saving techniques with reduced hands-on time are appreciated.

We report here the direct transposase-based library generation and nanopore sequencing of ssDNA as a convenient and versatile tool for characterization of AAV packaged DNA and ssDNA in general. We present proof for direct ssDNA sequencing by use of bacteriophage M13mp18 ssDNA as control (Figure 1 A). Use of a transposases for library generation was originally designed for dsDNA tagmentation (Nextera library preparation using Tn*5* transposase) and sequencing on the Illumina platform (31) and it was adapted for direct dsDNA sequencing on nanopores by Oxford Nanopore Technologies with MuA transposase (23, 32). MuA forms a homotetrameric synaptic complex around paired phage Mu genome ends and then catalyzes strand transfer, leaving behind nicks that act as replication primers in the wild type (33). A transposome consisting of MuA and end substrates (mini-Mu) is sufficient for *in vitro* transposition (34) and it shows only a slight target DNA bias towards a 5′-CYSRG pentamer (35, 36). Hyperactive MuA variants with also low target bias have been reported (37, 38). However, we were unable to find previous reports of MuA (or other DDE transposase superfamily members) activity on ssDNA targets. Our data confirms the relatively low insertion bias of MuA on dsDNA (Figure 1 B), but not on M13 ssDNA, where hot spots of insertions are seen, and activity is reduced. We observed three especially prominent hot spots of transposition sites for M13 ssDNA. Reads from M13 ssDNA exhibited lower sequence similarity compared to the reference than M13 dsDNA reads.

### Transposase activity and substrate

Our analysis showed that hot spots of transposase insertion sites can be in part explained by MuA action on (transient) hairpin loops within the ssDNA target. To fit into the MuA target binding pocket, hairpins require a stem of at least 23 – 25 nt (33, 36), however we saw also smaller hairpins as possible transposase targets. Moreover, MuA exhibits increased activity on mismatched targets (39) and the extent to which mismatches are tolerated has not yet been investigated, impeding our efforts to predict the target structures. Nonetheless, as we found the corresponding pairs of long reads and short reads for a given transposase insertion point on the circular substrate, we conclude that hairpin loops are one mode of transposase action on ssDNA. Compared to the M13 ssDNA sample, the AAV samples exhibited an overall relatively even distribution of transposition sites on both strands with a remarkable symmetry. It thereby closely resembles the dsDNA M13 sample. This effect is probably due to hybridization of two ssDNA genomes of AAV to one dsDNA, which we also observed in agarose gel electrophoresis. In the period between publication of our preprint and preparation of the manuscript for journal submission, another group confirmed the feasibility of our technique by tagmentation of hybridized AAV genomes and Illumina dye sequencing (40). Overlaid, we further find hot spots of transposition sites on one strand only. This is probably a different effect of transposase action on ssDNA, as seen for the M13 ssDNA. Given that hot spots lay on one strand only, we deduce that the effect could be in part sequence-specific in addition to conformation-specific, because of the inherent symmetry of hairpins when both strands are present. Further work will be required to completely elucidate the MuA transposase action on ssDNA targets.

### ITR sequencing and SNVs

Albeit the given insertion bias of the transposon on ssDNA, nanopore long reads compensated for this and still enabled full coverage of the recombinant AAV genome. We achieved a 356,009-fold coverage (run 2), which also enabled investigation of single-nucleotide variants despite the lower base accuracy of nanopore sequencing compared to other NGS methods. The sudden halving of coverage within the ITRs can be explained by a strong reduction of base-quality in the ITR regions, either stemming from difficulties of the helicase with the strong ITR secondary structure, or as a result of back-folding of ITRs after passing through the transmembrane pore. Furthermore, ITR internal hairpins can be present in two possible states, FLIP and FLOP (41), which adds to the reduction in coverage. A previous AAV NGS study found SNVs within the rAAV genome, mostly located within one region in the coding sequence and both ITRs (12). Regarding the ITRs, we found variants that locate in the ITR B-C hairpins (according to the ITR naming convention) as well. However, these are again attributable to the two possible states of ITRs which arise from AAV genome replication: FLIP and FLOP, where FLIP ITRs harbor the inverse complement C′-B′ hairpins compared to FLOP ITRs while the rest of the ITR sequence stays the same. Our producer plasmid pITR encodes FLOP ITRs on both sides and FLIP-specific SNVs appeared with a 20% frequency. We note that only those ITR conformation-specific (FLIP) SNVs were called, that lie in the outer arms of the respective hairpins, and we are so far unable to explain this finding. Also, SNVs within the coding sequence with frequencies up to 30% were observed. The same workflow resulted in only two called variants for the bacteriophage M13 sample as a control. Given that variant calling from nanopore data is a relatively recent technique, we suggest re-cloning of AAV DNA and Sanger sequencing of individual clones to confirm these high SNV abundancies. Nonetheless, our finding is overall in agreement with the previous study (12) which found SNVs with an abundance up to 15%. The locally high SNV abundance raises the question where these variants come from and what their implication for vector quality is. We find it unlikely that these SNVs are already present on the producer plasmid level, as this would render cloning in *E. coli* in general impractical to impossible and is also not in accordance with our frequent Sanger sequencing of pITR plasmids after cloning steps. Much rather, a somewhat error-prone AAV genome replication during virus production may be responsible and it would be very interesting to compare different producer cell lines (different mammalian, insect and yeast) and wild-type AAV under this aspect.

### Methylation of the packaged AAV genome

Nanopore sequencing also offered us the convenient opportunity to investigate CpG methylations from raw reads. In a previous study, laborious bisulfite PCR sequencing for packaged AAV2 wild type genomes showed little to no methylation with a maximum share of 1.7% methylated CpG dinucleotides, but revealed hypermethylation of integrated genomes (42). We used recently published deep learning tools to investigate CpG methylations from nanopore raw data, but we did not observe significant methylation above an unmethylated reference. The finding supports the previous study, which used AAV wild-type and highlights the similarity of wild-type and recombinant genome replication.

### Sample preparation and library preparation

As we present a direct sequencing method, the sample input is higher compared to other NGS methods. We find that, in case of AAV, extracted DNA from 10^13^ DNase I-resistant particles is enough for about five sequencing reactions. We also saw that a critical step in sample preparation is the Benzonase digest of the producer cell lysate. When performed for one hour with the given concentration, no fragments beyond the AAV packaging limit of about 5 kb are seen (Figure 2). Digestion for only 30 minutes on the other hand led to emergence of longer reads in small proportions (Supplementary Figure S8) and since we performed virus precipitation and antibody-based affinity chromatography for sample preparation, we attribute these to overlong fragments protruding the capsid and otherwise capsid-associated DNA. In the future, this incomplete (or omitted) digest could be used as a method to investigate rAAV genome replication and packaging intermediates directly. Also, there does not seem to be a linear correlation between sample DNA input and total read output, and we recommend using samples of OD_260, 10mm_ = 0.8 or higher for library preparation. Furthermore, the incubation time of the transposase can be optimized to yield longer fragments. In multiplexing we observed overspill and a lower read count, which may also be attributed to the differences in library preparation for multiplexing. We therefore recommend non-multiplexed sequencing, possibly on single-use flow cells (Flongle), for quality control settings.

### Sequencing of the packaging signal sequences

In our opinion, the relatively high amount of sample input is rewarded by the increased information content in long reads from direct sequencing. We were intrigued to see that 76% of AAV reads had a 3′-ITR sequence which is the predominant packaging initiation signal. Since the packaging direction is 3′ to 5′ (43, 44) and the sequencing direction is 5′ to 3′, reasons for reads terminating 3′ without an ITR could be two-fold: Either the corresponding fragment was double-fragmented by transposases prior to sequencing, or packaging initiated without an ITR. Given that all investigated human genomic sequences had no 3′-ITR, we find that both hypotheses remain valid, and no definite conclusion can be made from the given data. We can, however, safely say that the extent to which reads represent double-fragmented sequences was below 24% in our experimental setup. Another interesting analysis would be to investigate genomes that are incompletely packaged after packaging is initiated by the ITRs (truncated genomes). Since the sequencing direction opposes the packaging direction, this analysis is not possible with the provided method and, for example, 3′ adapter ligation, complementary strand synthesis and nanopore sequencing of the newly synthesized strand would be required.

As the AAV genome 3′-packaging signal is so conveniently accessible, we took the opportunity to search for 3′ p5 promoter sequences. The p5 promoter has been described to be a secondary packaging and genome replication signal (5, 45) and the corresponding activity was mapped to nucleotide positions 250 to 304 of the AAV serotype 2 reference genome with the terminal resolution site between positions 287 and 288 (46). Our data directly show that reads which terminated within the p5 promoter of the pRepCap plasmid, did so preferably at position +12 upstream of the TATA-box (equivalent to position 273 of the AAV serotype 2 reference genome). We reason that this 14 to 15 nt discrepancy between read-end position and the proposed terminal resolution site is probably an artefact of nanopore sequencing, in which the bases closest to the nucleic acid fragments 3′-end cannot be sequenced. We already observed this in our read-start and read-end analysis on M13 ssDNA, where the read-ends were shifted slightly by about 11 nt. Since the effect might also be sequence dependent and no systematic studies on this topic are available to us, we conclude that our data on the p5 promoter is in good agreement with previous studies. However, we also found previously undescribed recombination between p5 and ITR sequences. In these, the complete p5 promoter sequence was directly followed by the ITR, starting with the D-sequence. This means that the terminal resolution sites in the ITR and the p5 promoter are likely not involved in this recombination and it is unclear from our data how these sequences emerge. Although the overall proportion of these reads was small in our study, since p5 and ITR sequences were associated with *rep* and *cap* genes, we find that these sequences might contribute significantly to already observed packaging, persistence and expression of associated *cap* and also *rep* sequences from recombinant vectors (7, 10, 30, 47).

### Contaminant sequence abundancies and comparison to qPCR

We then wondered if our method also allows for quantification of contaminants in AAV vector stocks and we assigned reads to single entries of our reference database by BLASTn bit score as a first test and compared the results to our qPCR measurements. In general, the obtained share of contaminants was in the same range between the two methods. However, we found that the result depended also on the chosen read length threshold and in part also on the sequence numbering of the plasmid references. One source of deviation are long reads, that span more than one qPCR target. A distorted result is expected here, because of the unique read assignments which assigns one read to only the most prominent hit in the genome database and cannot assign possible fusions to both parental sequence sources. We tested a more advanced data evaluation technique with *in silico* read fragmentation and subsequent binning to a database reduced to the qPCR targets, but again, binning results were offset for *bla* and *rep* sequences. This hints on another source of error which might arise from the apparent transposase insertion bias we observed for ssDNA targets but might on the other hand also come from the qPCR where a primer set dependent bias was observed for AAV samples (48). Moreover, qPCR imposes a systematic error because human genomic sequences cannot be assessed. Normalization of reads by their respective read starts could level the transposase-induced error and might help to overcome the observed discrepancy to yield a quantitative method. Nonetheless, we find that qualitative statements on contaminations are already feasible. Concerning the length thresholding of reads for the BLAST analysis, we find that a threshold of >1,000 nt represents a good trade-off between depth of analysis, computing time and the emergence of double-fragmented sequences.

### Molecular state of packaged AAV genomes and ITR correction

To further characterize AAV packaged DNA, a %GC *vs*. read length plot proved to be a very convenient visualization for our nanopore data, as both parameters are computable quantities for uniquely fragmented sequences. When we plotted individual BLAST bins, a diverse picture emerged. First, read length histogram analyses allowed us to estimate that more than 99% of the rAAV genome sequences are of the expected size. Even though each read represents a fragmented genome, we find this conclusion is feasible, because both strands are equally likely packaged and independently sequenced, so that the non-strand-specific coverage in total is roughly constant along the genome length (evident in Figure 1 E). A simulation confirmed that we obtained mostly unique fragmentations, albeit with an observable data shift, probably due to incomplete sequencing of the 3′ ITR.

Second, further analysis showed that genome head-to-tail fusions are packaged in the capsid to a larger extent (Figure 2 B). The postulated primary mechanism for AAV genome replication suggests resolution of head-to-head and tail-to-tail fusions as rolling hairpin replication intermediates (44, 49), but we never observed such fusions. Head-to-tail fusions on the other hand could be the result of rolling circle replication, which requires a circular template. Circularized AAV genomes have long been described to occur *in vivo* as monomers or as head-to-tail fusions (50, 51). Further studies showed that these circular forms have a single junction ITR with a 5′ and a 3′ D-sequence (52) and that ssDNA genomes, which are subsequently packaged, can be replicated from this form by rolling circle replication (53). Compliant with those findings, fusions observed by us also had two D-sequences at the junction ITR. A further implication of the rolling circle replication model as a reservoir for ssDNA genomes is that the 3′ ITR sequence has to be repaired from the 5′ ITR as a template prior to genome packaging (53). This process can correct defective ITR sequences and ITR correction was observed even before circular intermediates were discovered (54). In our recombinant system, the plasmid carrying the GOI (pITR) encodes one ITR with a 11 bp deletion seen also by other groups (25), however we find now that this deletion is mostly absent in the ITRs of packaged genomes (except in the junction ITRs of genome-backbone fusions), which again hints on packaging from rolling circle amplified genomes. While the ITR “correction” mechanism in theory has equal probability to yield two ITRs with the deletion on the one hand and two correct ITRs on the other hand, it is not clear where the preference for correct ITRs in coming from. Given that fusions have correct ITRs hints on ITR correction as a preliminary requirement prior to genome amplification.

Notably, the mapping quality in the junction ITRs is low and it is not clear, whether this is a result of sequencing artifacts due to the strong secondary structures or due to error-prone recombination at this position. Given that the mapping quality always drops 3′ of the B-C hairpins, independent of the genome orientation, the former explanation seems likely, although error prone circularization has also been observed (50). Possible explanations for the combined findings of dominantly monomer genomes on the one hand and the observation of head-to-tail fusions on the other hand are threefold: Either i) rolling circle replication rather than rolling hairpin replication might be the predominant replication scheme in our setting, possibly enforced by the erroneous ITR, or ii) after gene correction through rolling circle replication, replication could also proceed mostly by rolling hairpin replication, provided that genome monomer excision is more efficient in this replication scheme. Equally possible is that iii) observed fusions occur because their respective rolling circle templates harbor erroneous ITRs due to erroneous recombination and the resulting concatemers cannot be resolved to monomers following genome amplification, whereas the majority of genomes stem from templates with correct ITRs, where resolution is efficient. Without further investigation, for example on cellular rAAV unpackaged DNA, no definite conclusion is possible.

### Packaging of pITR backbone and other contaminant sequences

We also observed AAV genome-plasmid backbone fusions with identity between the fusions and the pITR plasmid. In these, the junction ITR still harbored the 11 nt deletion and we observed a higher mapping quality throughout the junction. It is possible that nanopore sequencing is facilitated by ITRs with the deletion, which does not form as-strong secondary structures as wild type ITRs. These fusions furthermore terminated mostly in an ITR and we observed them in both polarities (GOI located 3′ and backbone located 3′), so that we suggest that they might also be actively replicated and not packaged directly from the producer plasmid. Observations of solely bacterial backbone sequences with 3′ terminal ITRs within the pITR bin add to this suggestion. ITRs were found also 3′ terminal of *rep* and *cap* as well as adenoviral helper sequences providing direct confirmation of previously observed non-homologous recombination between ITRs and helper sequences (47, 55) and warrants efforts for their elimination (56), although the overall quantity of these reads was low in our setting.

In contrast, human genomic sequences did not harbor 3′ ITR sequences. Instead, they were found to be randomly packaged and of exponentially decaying size distribution, which hints at the involvement of enzymatic digestion. This raises the question, where these fragments originate from. We find it unlikely that the host cell genome is highly fragmented during virus production, however, host cell DNA is treated with Benzonase nuclease during virus purification. We wonder whether residual helicase activity of the AAV replicases during Benzonase treatment of producer cell lysate is responsible for randomly packaged host cell DNA. Presumably, host cell DNA could be randomly packaged from free ends during purification and is then cut at random timepoints at the capsid surface, yielding the decaying size distribution. Work towards a specific replicase inhibitor that can be added during virus purification might be a chance to further improve vector quality.

In conclusion, we present here unprecedented deep nanopore sequencing of packaged ssDNA in recombinant AAV with the possibility to expand the application range to other single-stranded viruses or bacteriophages, as demonstrated for bacteriophage M13 circular ssDNA. The technique dramatically simplifies sample preparation and reduces turnover times compared to other NGS characterization methods for ssDNA and the information content of the result increases due to long reads. Deep and direct nanopore sequencing provided us with direct confirmation of several fundamental research discoveries on AAV biology: From statements on AAV genome primary and secondary packaging signals and genome orientation over the genome’s replication scheme and single-nucleotide variants to assessments on recombination events. Furthermore, direct sequencing allowed for qualitative statements on DNA contaminations in AAV vector stocks. The present study highlights the necessity to further understand the AAV basic biology to gain high-transducing vectors with homogeneous payloads for gene therapy applications. Analytical procedures must keep pace with new diagnostic developments, and we foresee that quantitative PCR will lose its status as the gold standard, as unbiased next-generation sequencing protocols become cheaper, easier and readily available.

## AVAILABILITY

Custom Python scripts are available in the GitHub repository (https://github.com/MarkusHaak/Radukic_Brandt_2020).

## ACCESSION NUMBERS

Sequencing data have been deposited with the Sequence Read Archive (SRA) under accession number PRJNA610225. A reviewer link is provided: https://dataview.ncbi.nlm.nih.gov/object/PRJNA610225?reviewer=4m40cdtp0gscs7k5mdqfnprbfu

## SUPPLEMENTARY DATA

Supplementary Data are available online.

