## Supplementary Data for "Nanopore sequencing of native adeno-associated virus (AAV) single-stranded DNA using a transposase-based rapid protocol"

### 1 Methods

#### 1.1 Production of M13KO7 helper phage

*E. coli* ER2738 from an over-night culture were used to inoculate lysogeny broth (LB) supplemented with ampicillin (100 µg/ml) and tetracycline (25 µg/ml)=to $OD600_{10mm}$ = 0.1 and grown to $OD600_{10mm}$ = 0.5 at 37 °C in Erlenmyer flasks on an orbital shaker. Next, the culture was infected with 4 × $10^9$ pfu/ml M13KO7 (New England Biolabs) helper phage and further incubated for 1.5 hours. Kanamycin was added to a final concentration of 70 µg/ml and, after further four hours of incubation, the culture supernatant containing the phages was separated from cells by centrifugation. Phage ssDNA was prepared from the supernatant by the QIAprep Spin M13 Kit (Qiagen) as per the manufacturer's instructions. 6.2 µg ssDNA were obtained from a 3 ml preparation.

#### 1.2 qPCR assay

We performed all qPCR measurements with primers and reagents according to the following tables. All dilution series for standard curves and primer dilutions were performed with sterile filtered Millipore MilliQ water in presence of 0.05% Pluronic F68. All pipetting was done with Sarstedt Biosphere low retention filter tips. Final primer concentration was 125 nM. The reaction volume was 20 µl in 96 well Sarstedt Lightcycler plates. Quantification cycles were determined with the Roche $2^{nd}$ derivative Max algorithm within the LightCycler 480 software, release 1.5.0 SP4, version 1.5.0.39.

All measurements were performed in technical duplicates. Sample DNA was extracted as described in the methods section and further diluted for the assay with MilliQ water containing 0.05% Pluronic F68. Nucleic acid quantification of AAV producer plasmids for standard curve preparation was performed spectroscopically with a Nanodrop 2000c. The ratio OD260/OD280 was between 1.85 and 1.88.

Table 1: Primer sets used for qPCR measurement.

| **Target** | **Primer sequences** | **Amplicon length** | **qPCR program** | **Primer efficiency** | **Linear range** | **Limit of detection** |
|---|---|---|---|---|---|---|
| CMV promoter | 5‘-GGGACTTTCCTACTTGGCA<br>5‘-GGCGGAGTTGTTACGACA | 200 bp | A | 1.84 | $10^3$ – $10^9$ per reaction | <$10^3$, * |
| AAV serotype 2 Rep | 5’-CGGAGAAGCAGTGGATCCA<br>5’-ATTTGGGACCGCGAGTTG | 76 bp | B | 1.82 | $10^3$ – $10^9$ per reaction | <$10^3$, * |
| Adenovirus gene E4 | 5’-ACTACGTCCGGCGTTCCAT<br>5’-GGAGTGCGCCGAGACAAC | 68 bp | A | 1.85 | $10^3$ – $10^9$ per reaction | <$10^3$, * |
| beta lactamase (Ampicillin resistance gene, *bla*) | 5’-CAACTTTATCCGCCTCCATC<br>5’-AAGCCATACCAAACGACGAG | 138 bp | A | 1.91 | $10^3$ – $10^9$ per reaction | <$10^3$, * |

* Standard error of regression method

Table 2: qPCR programs.

| **qPCR program** | **Instrument, Assay** | **Master Mix** | **Program** |
|---|---|---|---|
| A | Roche LightCycler 480II SYBR-Green type assay | Promega GoTaq qPCR Master Mix | a. 95 °C, 10 min.<br>b. 95 °C, 15 sec.<br>c. 60 °C, 1 min.<br>d. to b., 39x<br>e. 45 °C to 95 °C at 0.11 °C/sec |
| B | Roche LightCycler 480II SYBR-Green type assay | Promega GoTaq qPCR Master Mix | a. 95 °C, 10 min.<br>b. 95 °C, 15 sec.<br>c. 55 °C, 15 sec.<br>d. 60 °C, 1 min.<br>e. to b., 39x<br>f. 45 °C to 95 °C at 0.11 °C/sec |

Table 3: Standard curves for qPCR primer sets

| **Primer set** | **Fluorescence graph and melting curve** | **Standard curve** |
|---|---|---|
| CMV | 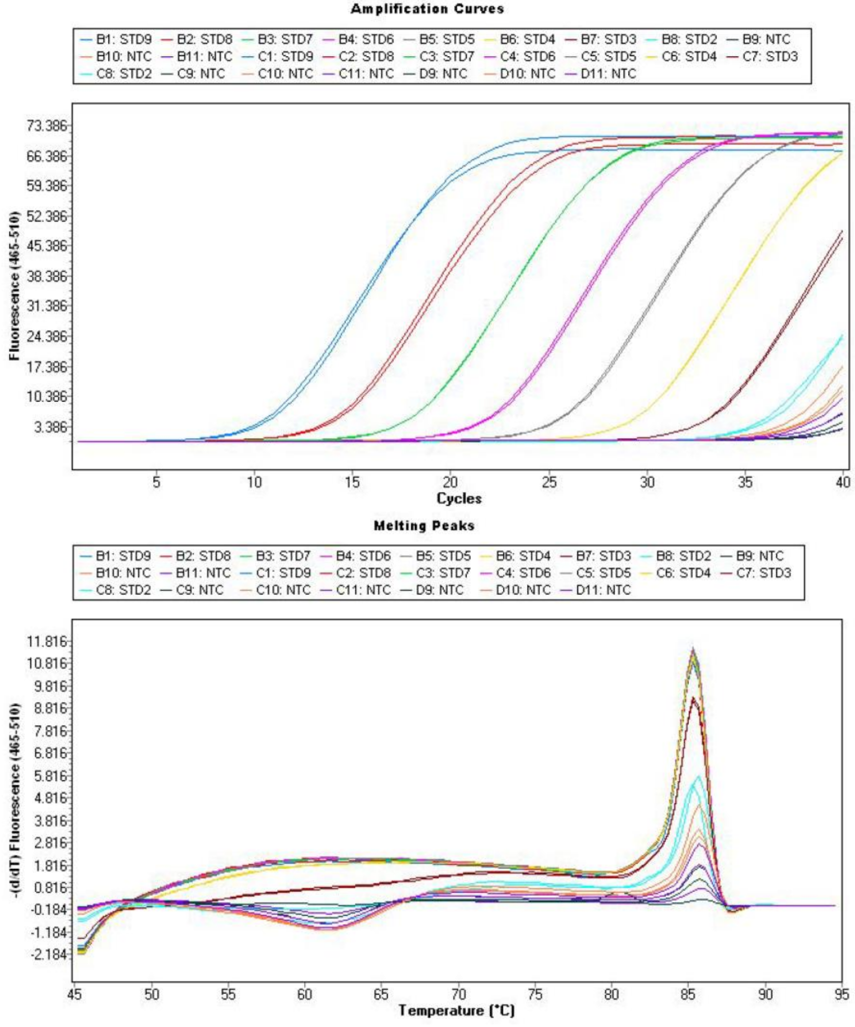<br> | 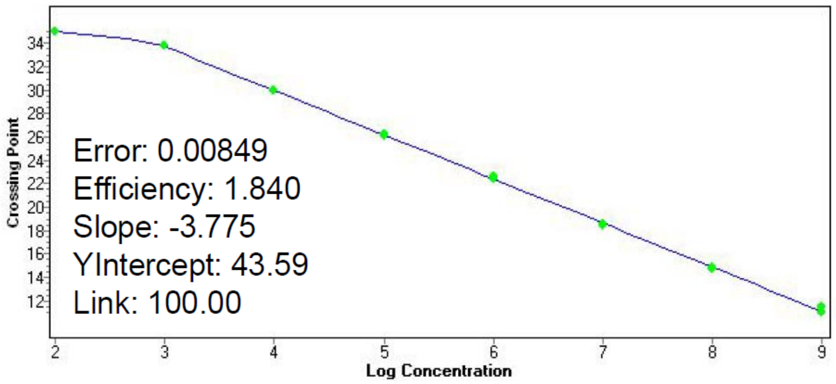<br> |

| | | |
|---|---|---|
| Rep | 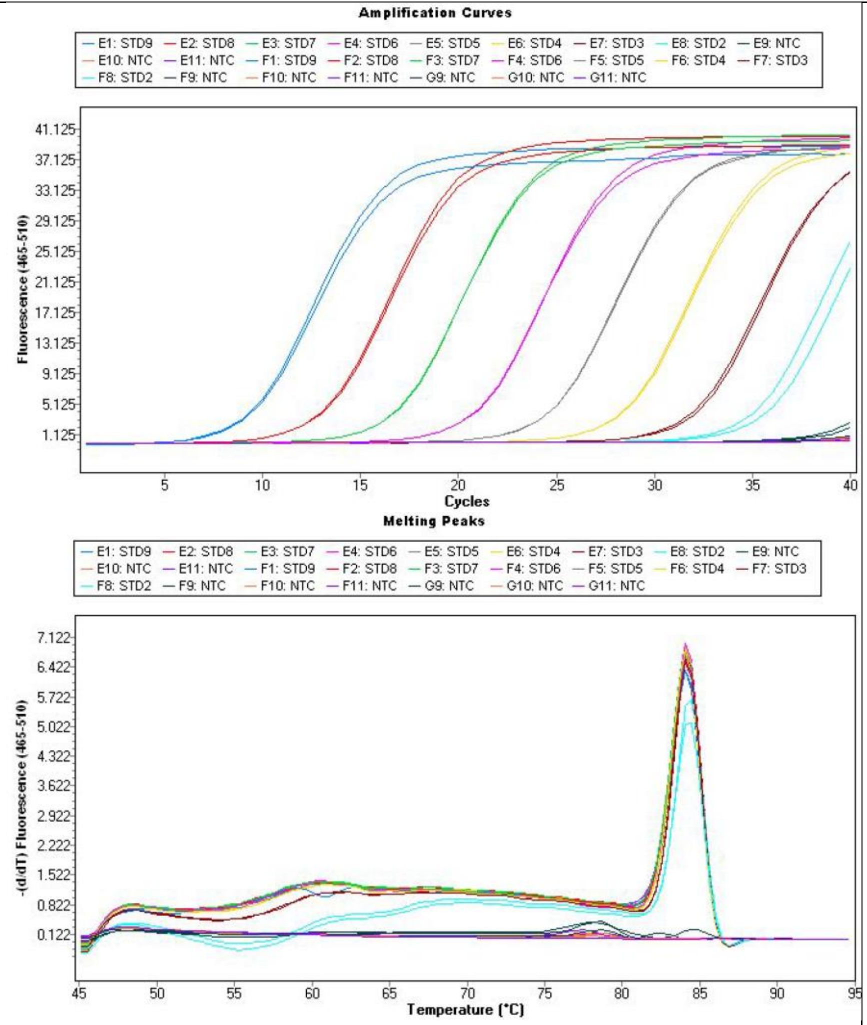Amplification Curves<br>E1: STD9 E2: STD8 E3: STD7 E4: STD6 E5: STD5 E6: STD4 E7: STD3 E8: STD2 E9: NTC<br>E10: NTC E11: NTC F1: STD9 F2: STD8 F3: STD7 F4: STD6 F5: STD5 F6: STD4 F7: STD3<br>F8: STD2 F9: NTC F10: NTC F11: NTC G9: NTC G10: NTC G11: NTC<br>Fluorescence (465-510)<br>Cycles<br>Melting Peaks<br>E1: STD9 E2: STD8 E3: STD7 E4: STD6 E5: STD5 E6: STD4 E7: STD3 E8: STD2 E9: NTC<br>E10: NTC E11: NTC F1: STD9 F2: STD8 F3: STD7 F4: STD6 F5: STD5 F6: STD4 F7: STD3<br>F8: STD2 F9: NTC F10: NTC F11: NTC G9: NTC G10: NTC G11: NTC<br>-(d/dT) Fluorescence (465-510)<br>Temperature (°C) | 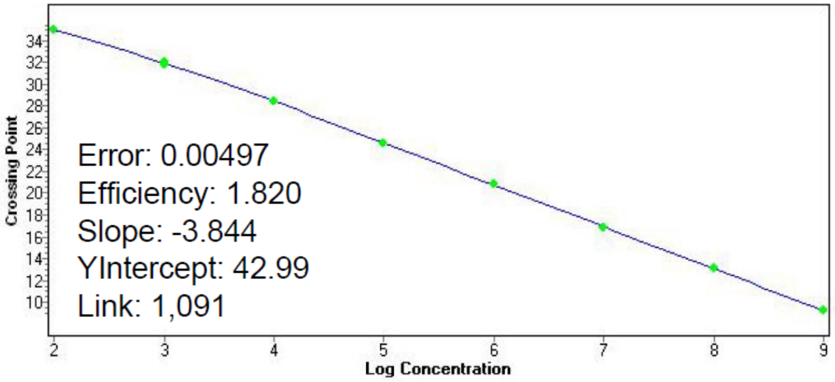Error: 0.00497<br>Efficiency: 1.820<br>Slope: -3.844<br>YIntercept: 42.99<br>Link: 1,091<br>Crossing Point<br>Log Concentration |

| | | |
|---|---|---|
| E4 | 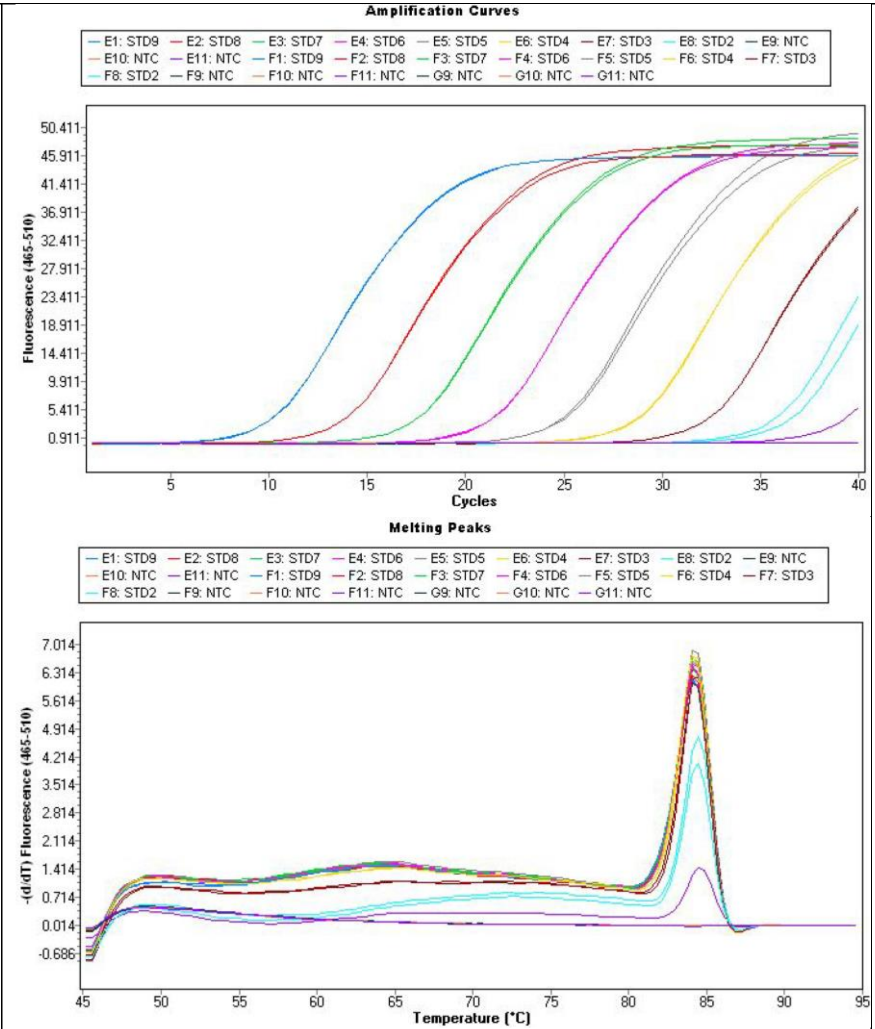Amplification Curves<br>E1: STD9 E2: STD8 E3: STD7 E4: STD6 E5: STD5 E6: STD4 E7: STD3 E8: STD2 E9: NTC<br>E10: NTC E11: NTC F1: STD9 F2: STD8 F3: STD7 F4: STD6 F5: STD5 F6: STD4 F7: STD3<br>F8: STD2 F9: NTC F10: NTC F11: NTC G9: NTC G10: NTC G11: NTC<br>Fluorescence (465-510)<br>Cycles<br>Melting Peaks<br>-(d/dT) Fluorescence (465-510)<br>Temperature (°C) | 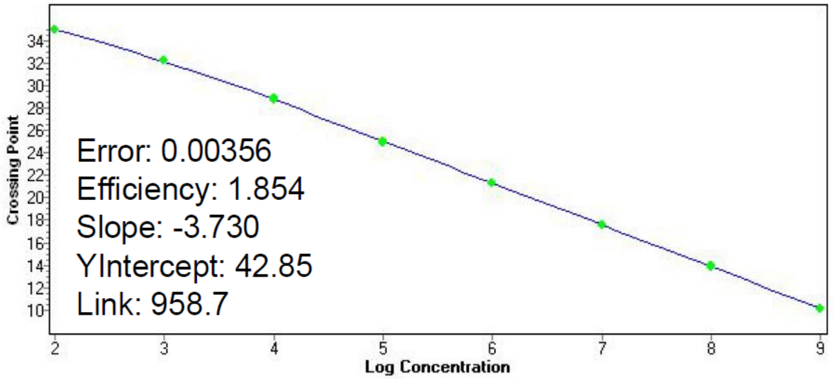Error: 0.00356<br>Efficiency: 1.854<br>Slope: -3.730<br>YIntercept: 42.85<br>Link: 958.7<br>Crossing Point<br>Log Concentration |

bla

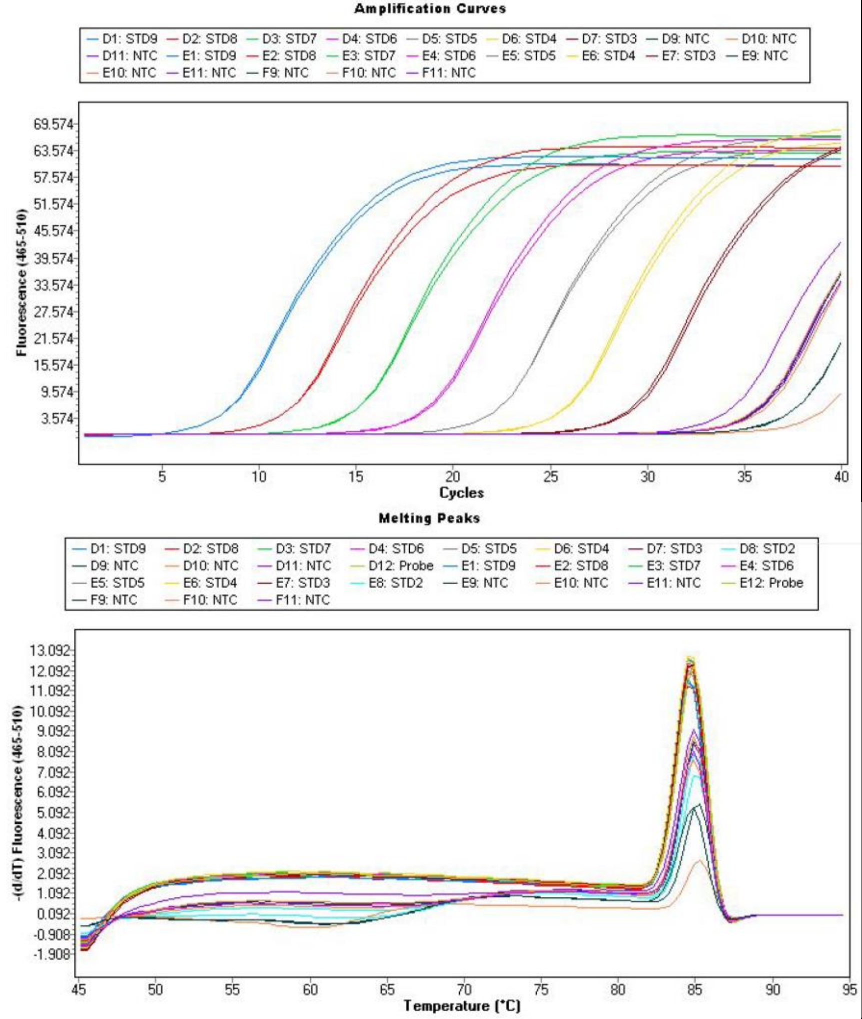
Amplification Curves
D1: STD9
D2: STD8
D3: STD7
D4: STD6
D5: STD5
D6: STD4
D7: STD3
D9: NTC
D10: NTC
D11: NTC
E1: STD9
E2: STD8
E3: STD7
E4: STD6
E5: STD5
E6: STD4
E7: STD3
E9: NTC
E10: NTC
E11: NTC
F9: NTC
F10: NTC
F11: NTC
Fluorescence (465-510)
Cycles
Melting Peaks
D8: STD2
D12: Probe
E8: STD2
E12: Probe
-(d/dT) Fluorescence (465-510)
Temperature (°C)

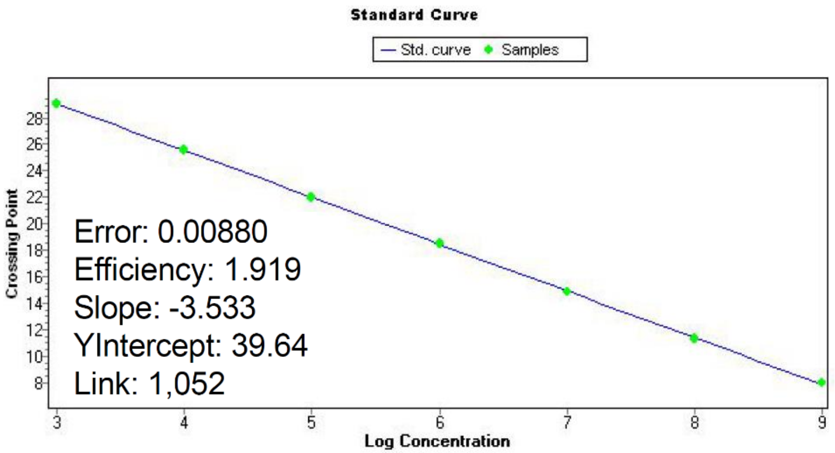
Standard Curve
Std. curve
Samples
Crossing Point
Log Concentration
Error: 0.00880
Efficiency: 1.919
Slope: -3.533
YIntercept: 39.64
Link: 1,052

#### 2 Script for the simulation of the transposase reaction

The following script was developed for and tested on GNU Octave 5.1.0. Sequences to be analysed can be put between the brackets in line 2. The script will output results to a graph and a text file in the user folder (under MS Windows).

```
1  #Input DNA Sequence
2  DNAseq="Put sequence of interest between brackets";
3  variants = length(DNAseq)-1;
4  r_m = ones (variants*2, 2);
5  
6  #Basic conversions
7  DNAseq( DNAseq == "G" )="S";
8  DNAseq( DNAseq == "C" )="S";
9  
10 #Calculate %GC upstream
11 for i = 1:variants
12   #length of DNA fragment
13   r_m([i], [1]) = variants-i+1;
14 
15   #%GC
16   r_m([i], [2]) = columns(strfind(substr(DNAseq, i+1), "S")) / r_m([i], [1]) * 100;
17 
18 endfor
19 
20 #Calculate %GC downstream
21 for i = variants+1:rows(r_m)
22   #lenght of DNA fragment (same as above)
23   r_m([i], [1]) = i-variants;
24 
25   #%GC
26   r_m([i], [2]) = columns(strfind(substr(fliplr(DNAseq), variants-i), "S")) /
27 r_m([i], [1]) * 100;
28 endfor
29 
30 save results.mat r_m
31 plotmatrix(r_m)
```

### 3 Transposase adapter sequences

The following transposase adapter sequences were used for the realignment of untrimmed reads in order to estimate transposition sites:

Samples M13mp18 ssDNA and M13KO7 ssDNA:

5′-GCTTGGGTGTTTAACCTTCAGGGAACAAACCAAGTTACGTGTTTTCGCATTTATCGTGAAACG CTTTCGCGTTTTTCGTGCGCCGCTTCA

Sample M13mp18 phagemid dsDNA:

5′-GCTTGGGTGTTTAACCAACTAGGCACAGCGAGTCTTGGTTGTTTTCGCATTTATCGTGAAAC GCTTTCGCGTTTTTCGTGCGCCGCTTCA

AAV sample 2 (run 2):

5’-GCTTGGGTGTTTAACCGTTTTCGCATTTATCGTGAAACGCTTTCGCGTTTTTCGTGCGCCGCTT CA

### 4 Supplementary Figures

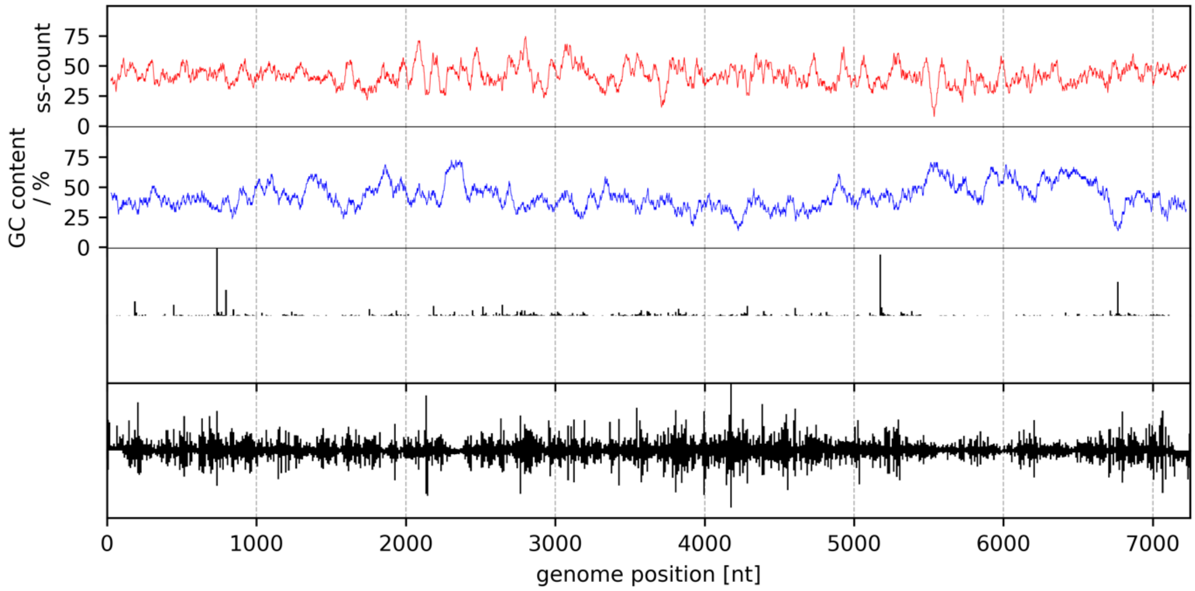

**Supplementary Figure S1:** Plus strand ss-count, GC content and relative transposase insertion sites of samples M13mp18 ssDNA (top) and M13mp18 dsDNA (bottom). The ss-count is based on 100 predicted DNA folding structures. Both ss-count and GC content are averaged over a moving window of 50 nt. The estimated read starts are binned in 15 nt bins and normalized to the maximal bin count.

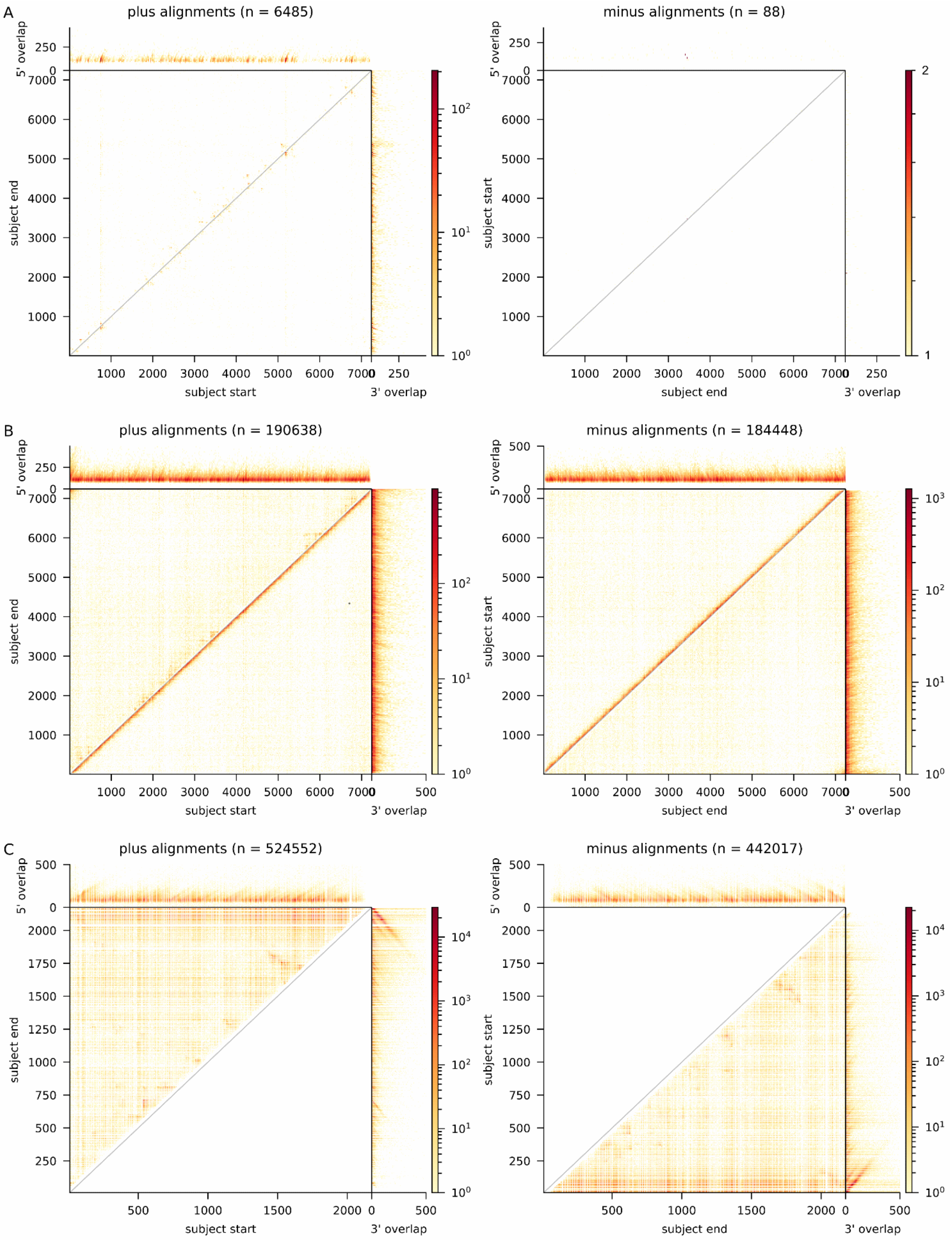

**Supplementary Figure S2:** Binned subject start and end positions for untrimmed reads mapped against ssM13 (A), dsM13 (B) and rAAV (C) genomes. Bins are sized 15 x 15 nt for ssM13 and dsM13 and 5 x 5 nt for rAAV. The length of unmapped stretches of reads is plotted as 5′ overhang and 3′ overhang, respectively. Read count per bin is scaled logarithmically.

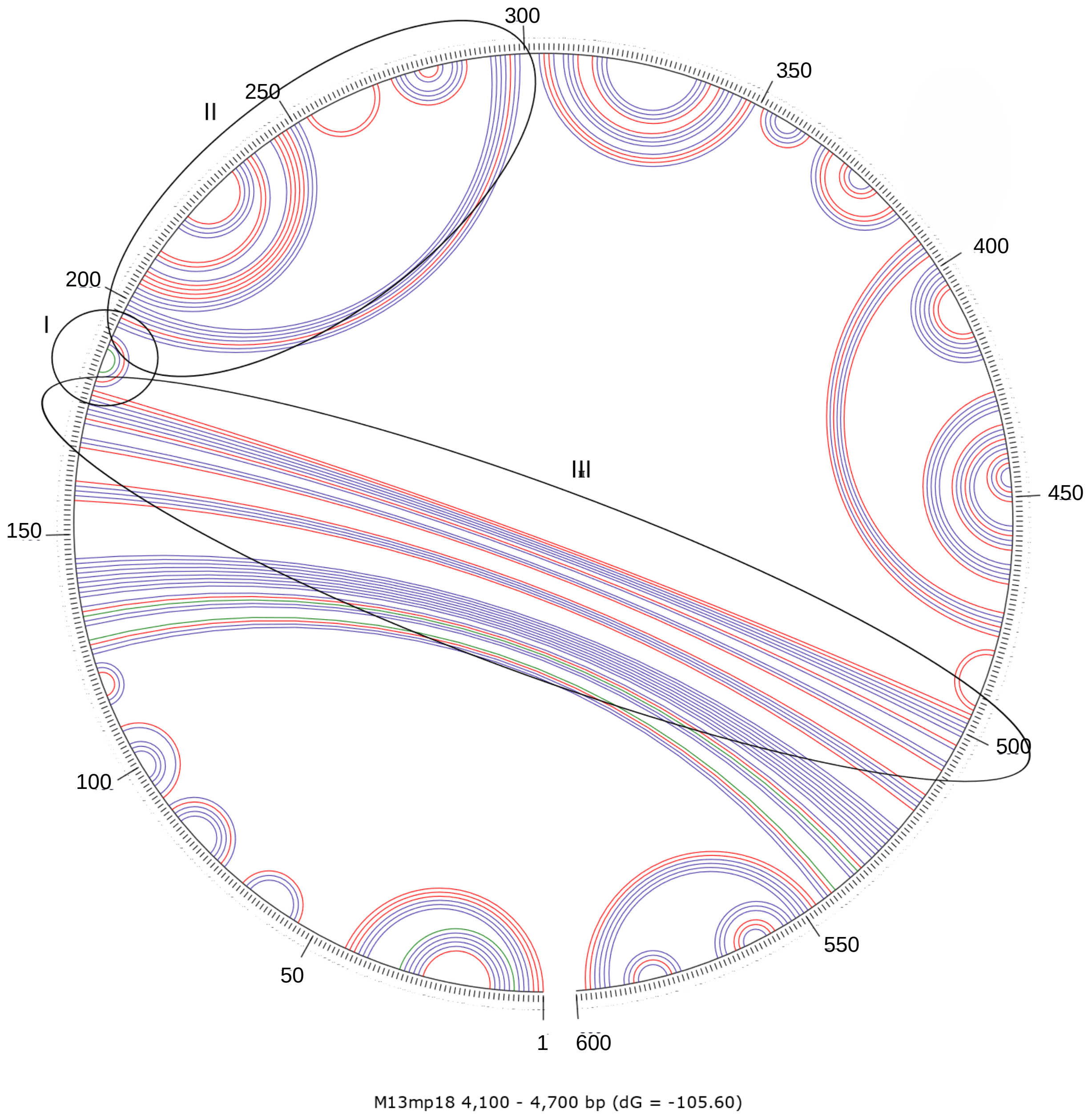

**Supplementary Figure S3:** Circular structure plot of predicted folding from base 4100 to 4700 of M13mp18 genomic DNA. Structure prediction carried out using mfold_util 4.7 with standard options. Possible hairpin loops serving as targets for transposase insertion are highlighted.

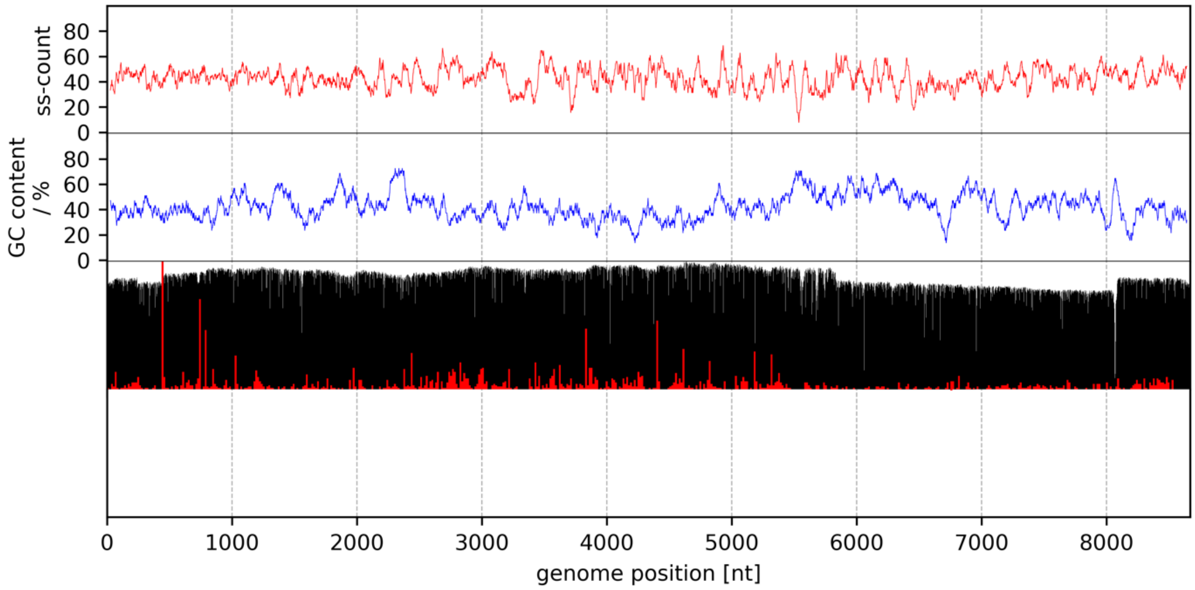

**Supplementary Figure S4:** Plus strand ss-count, GC content, relative coverage and relative transposase insertion sites of sample M13KO7 ssDNA. The ss-count is based on 100 predicted DNA folds. Both ss-count and GC content are averaged over a moving window of 50 nt. The estimated read starts are binned in 15 nt bins and normalized to the maximal bin count.

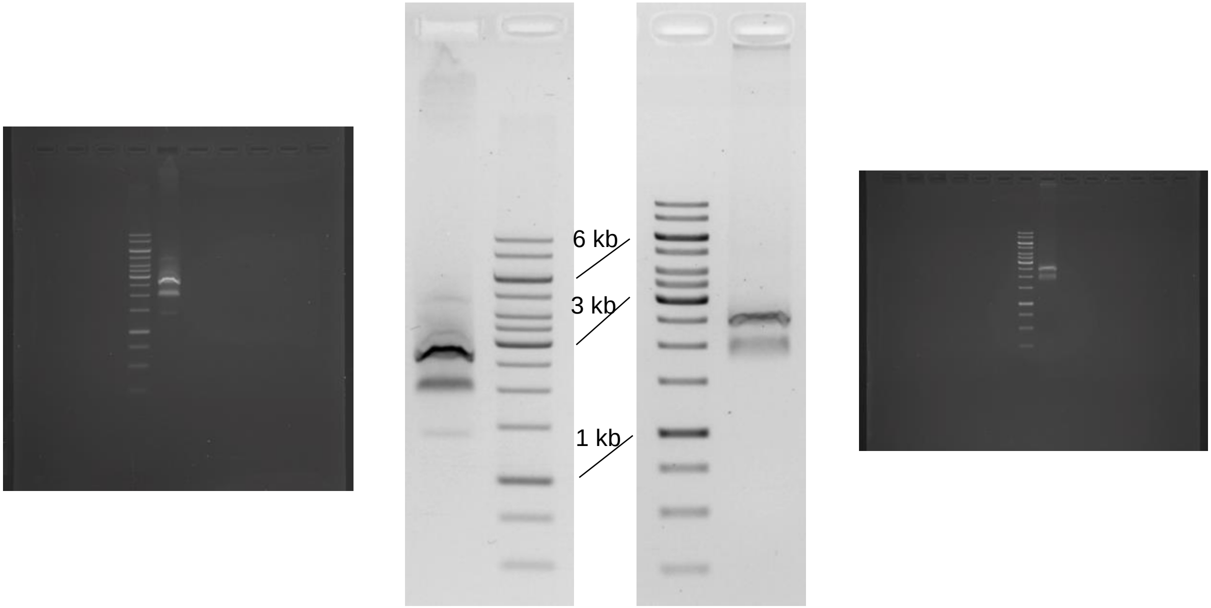

**Supplementary Figure S5:** Agarose gel electrophoresis of DNA prepared from rAAV sample 1 used for multiplexed sequencing (run 1). Both gels show a 5 µl sample from the same preparation before (left) and after a freeze-thaw cycle. The gel was 1% agarose in TAE buffer, run at 120 V for 50 min. Staining by SYBR Gold nucleic acid stain (Thermo). The marker was Gene Ruler 1 kb (Thermo). Gels displayed with inverted colors and spread histograms. Uncropped gels are given. The unfrozen sample shows three distinct bands. One band runs below the expected genome size of 2.2 kb. One runs right at the expected size and one runs between 2.5 and 3 kb. Upon freeze-thawing, the smallest fragment disappears, and aggregates are seen at the sample pocket, whereas the other bands are preserved. We assume that the smallest fragment resembles true single-stranded genomes, while the band of the expected size resembles two at least partly hybridized genomes. Higher order non-covalent multimers seem also to be present. Under the investigated conditions, a larger part of the sample seems to be in hybridized states.

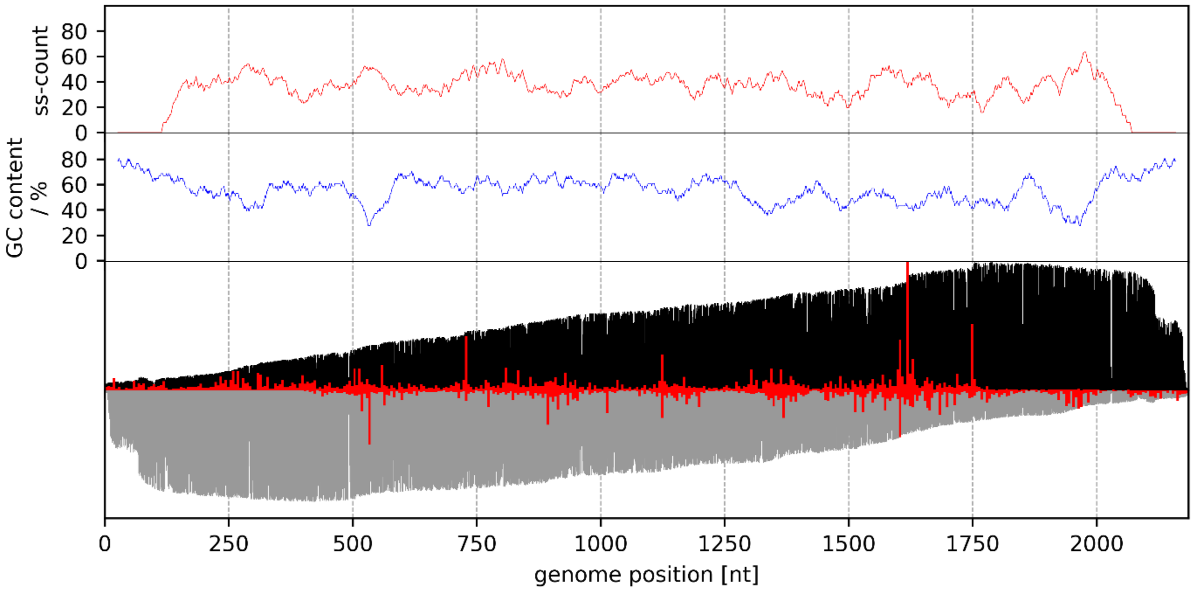

**Supplementary Figure S6:** Plus strand ss-count, GC content, relative coverage and relative transposition sites of sample rAAV (run 2). The ss-count is based on 100 predicted DNA folds. Both ss-count and GC content are averaged over a moving window of 50 nt. The estimated read starts are binned in 5 nt bins and normalized to the maximal bin count.

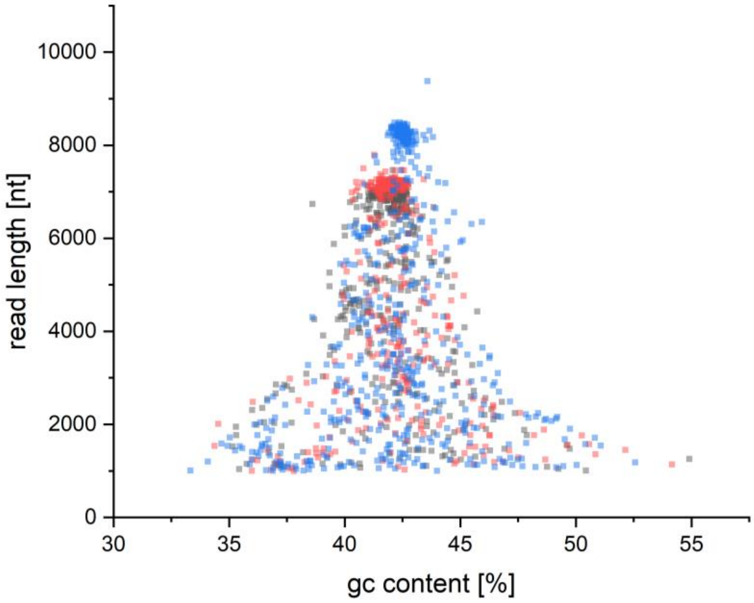

**Supplementary Figure S7:** GC content *versus* read length for a commercial M13mp18 dsDNA phagemid (grey, partially overlaid by red) and M13mp18 ssDNA (M13mp18, red), as well as M13 inhouse propagated helper phage (blue). Most reads are of similar length and GC content, indicating that an individual molecule is only fragmented once by the transposase. Conical tailing to minor extends hints on premature sequencing breakoffs and the prevalence of double-cut genomes.

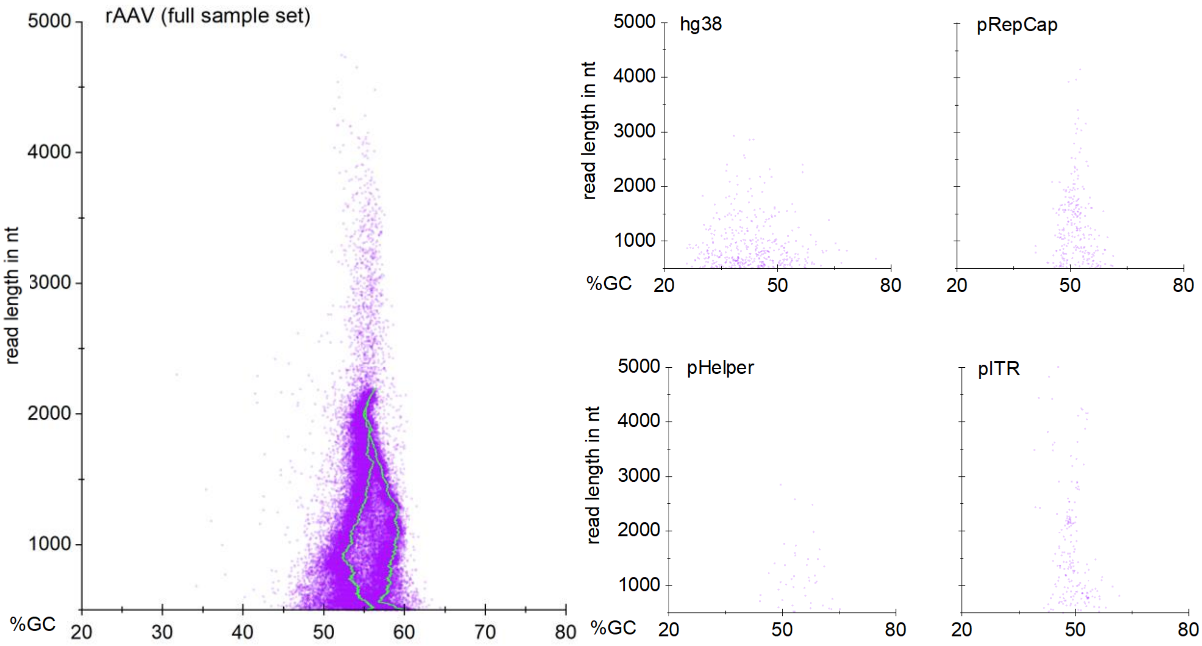

**Supplementary Figure S8:** GC content *versus* read length for the multiplexed AAV sequencing run, grouped by BLAST assignments to the reference library. One magenta dot represents on read. Of all 36239 reads that passed the quality threshold of >500 nt, 52 (0.14%) were longer than 5000 nt, 51 of which fell into the pITR bin. This finding highlights the importance of complete Benzonase digest prior to capsid disruption. Additionally, the data indicates that oversized reads likely stem from the pITR backbone instead of genome multimers or the other producer plasmids.

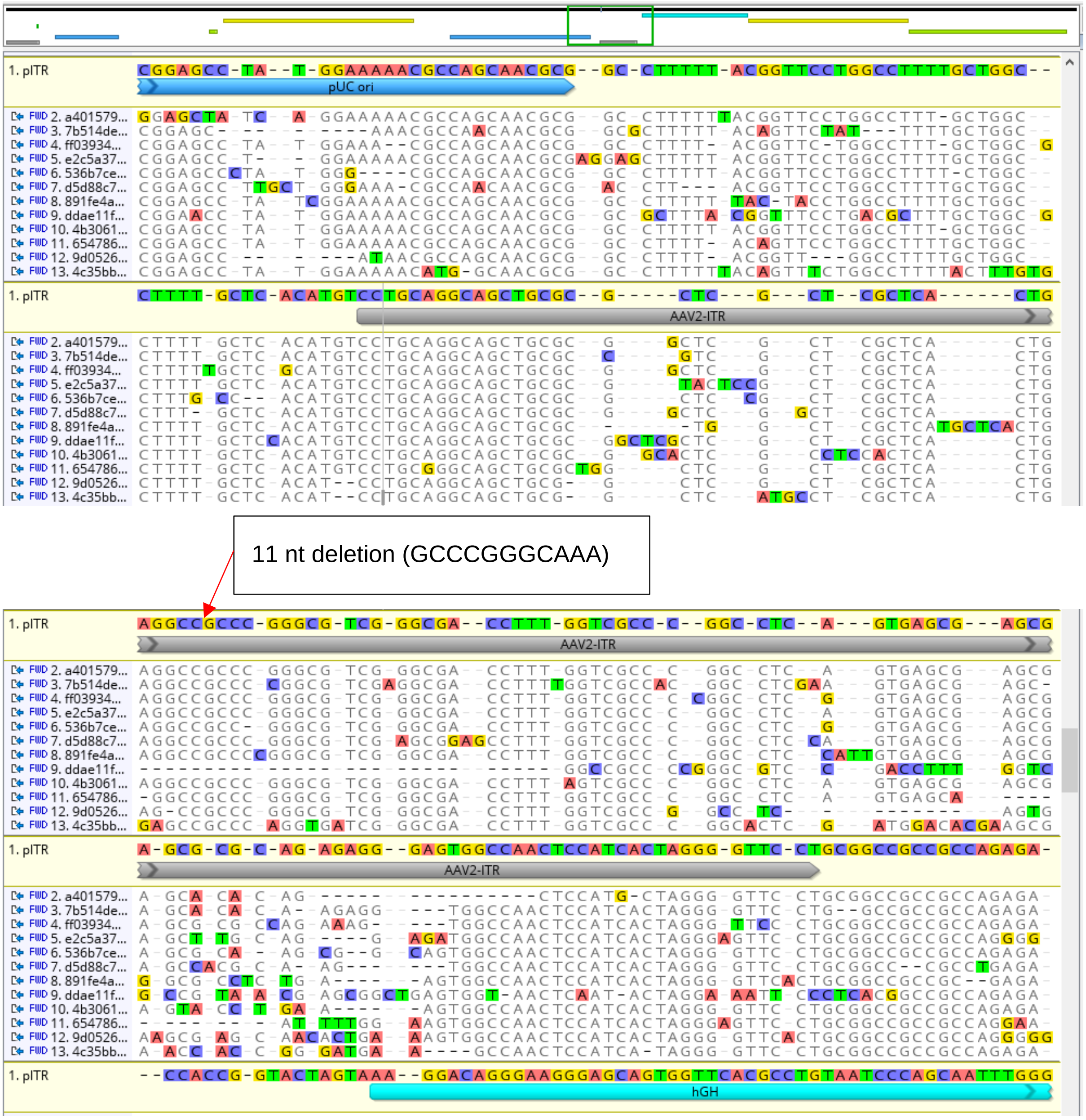

**Supplementary Figure S9:** 12 read examples aligned to the junction ITR between genome-backbone fusions are shown. Junction ITRs still harbour a 11 nt deletion and sequencing quality drops downstream of the ITR internal palindrome (12 read examples mapped to pITR).

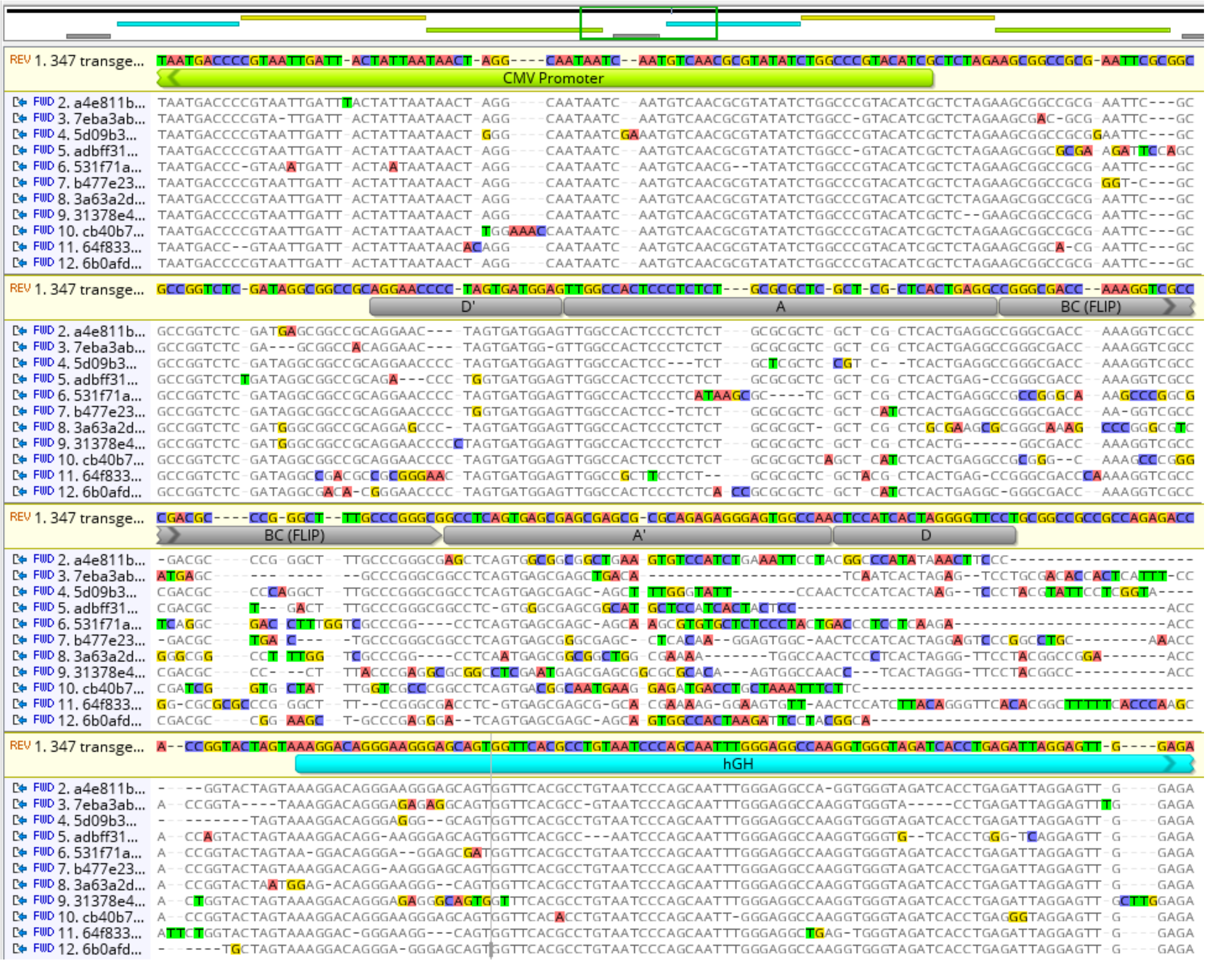

**Supplementary Figure S10:** Junction ITRs between genome-genome fusions have two D-sequences and sequencing quality drops downstream of the internal palindrome (BC). The 11 nt deletion present on producer plasmid level is absent in the genome-genome fusions (11 read examples mapped to *in silico* constructed genome-genome fusions).

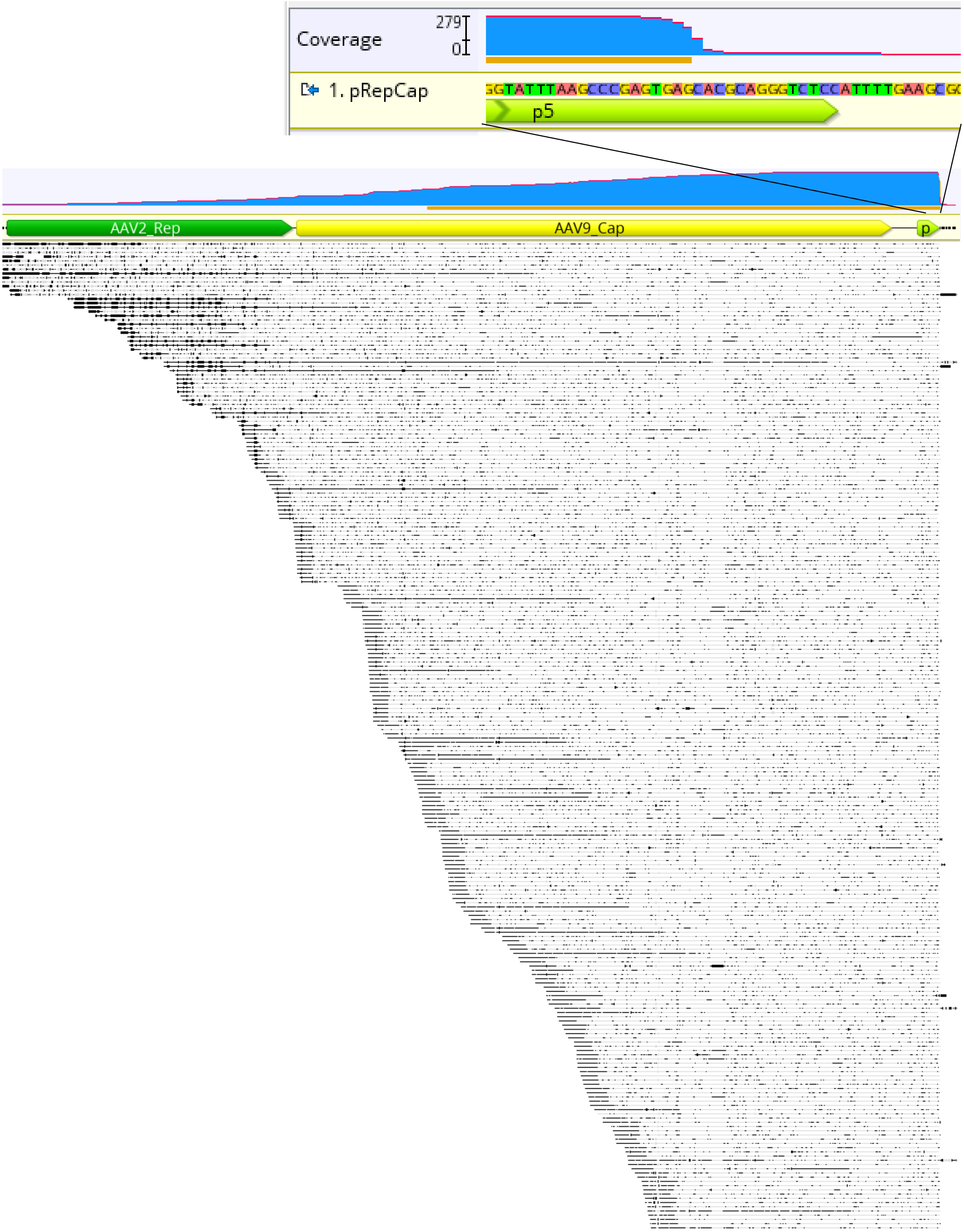

**Supplementary Figure S11:** p5 coverage drops sharply at position +12 of the promoter's TATA-box.

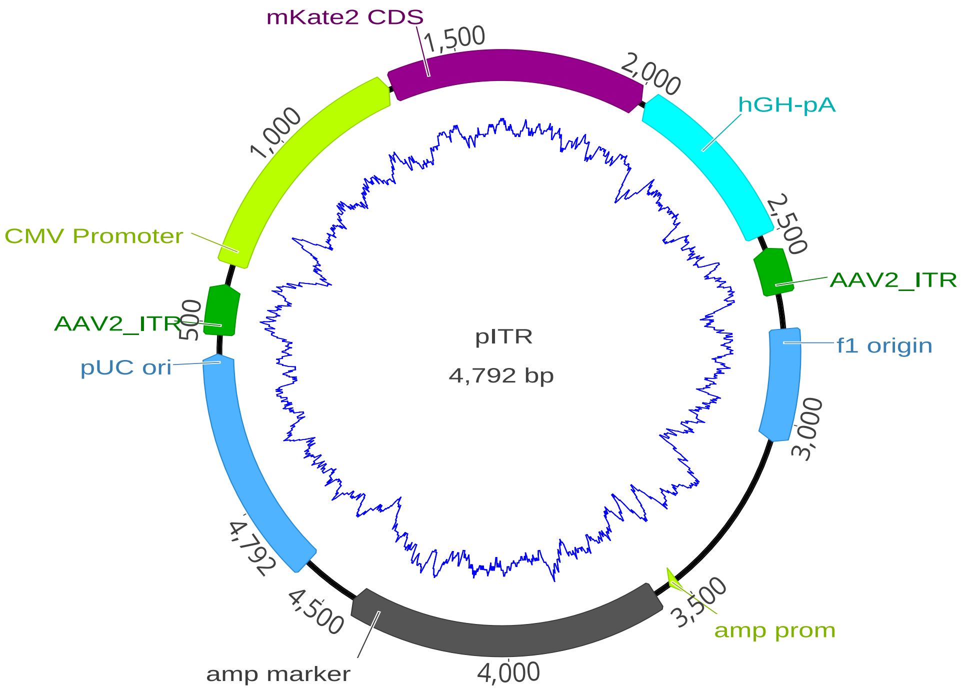

**Supplementary Figure S12:** Plasmid map of pITR (pUC19bb_ITR_EXS_CMV_mKate2_hGH-pA) with GC content graph. The right ITR (proximate to the hGH-pA) harbors a 11 nt deletion.

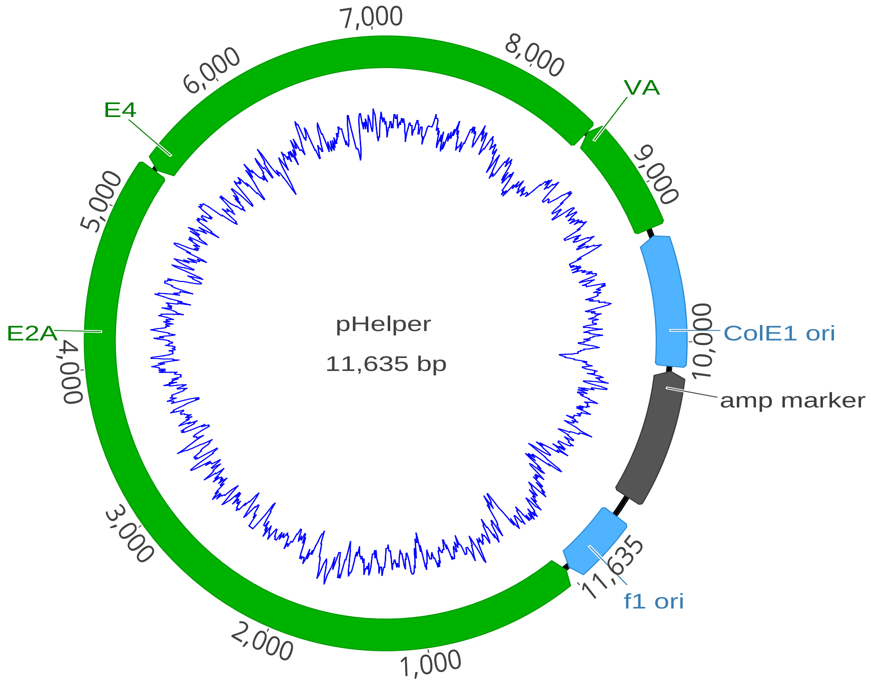

**Supplementary Figure S13:** Plasmid map of pHelper (Agilent Technologies) with GC content graph.

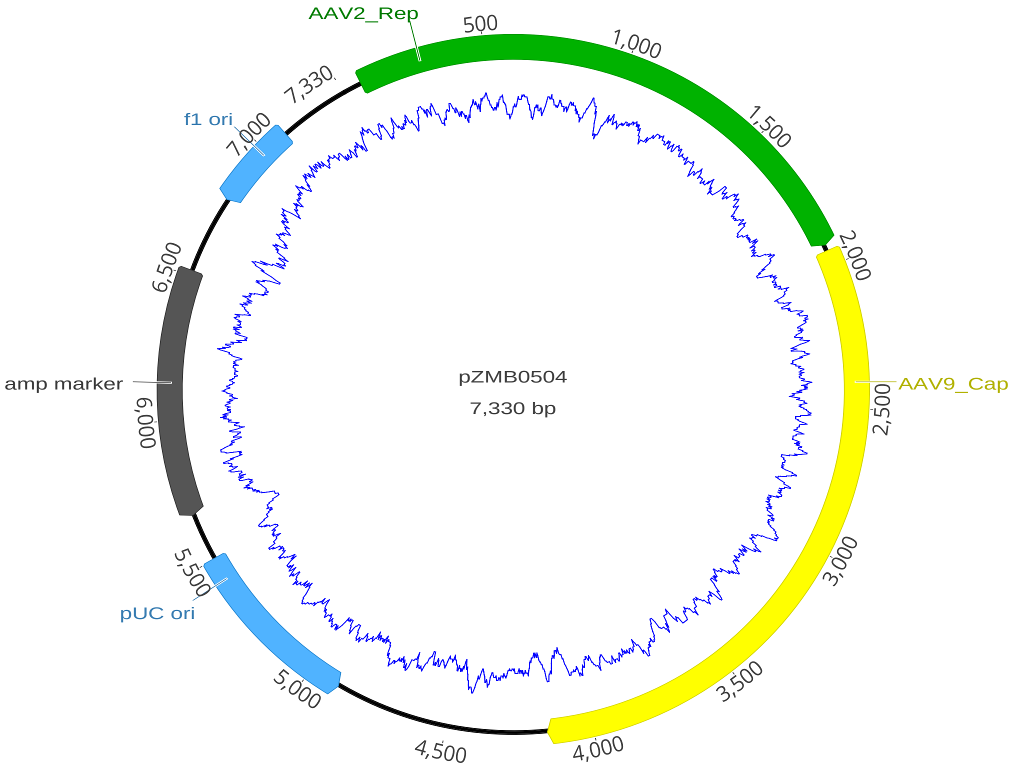

**Supplementary Figure S14:** Plasmid map of pRepCap (pRep2Cap9) with GC content graph.
